# Invariant natural killer T-cell and CD4^+^ T-cell derived IL22 is a regulator of epithelial-to-mesenchymal transition and extracellular matrix remodelling in perianal fistulas

**DOI:** 10.1101/2023.09.11.557122

**Authors:** Laura E Constable, Nusrat Iqbal, Domenico Cozzetto, Luca Csabai, Sulak Anandabaskaran, Tamas Korcsmaros, Ailsa Hart, Phillip J Tozer, Nick Powell

**Author notes:** Correspondence: Professor Nick Powell, MBChB, PhD, MRCP, Division of Digestive Disease, Department of Metabolism Digestion and Reproduction, Faculty of Medicine, Imperial College London, Du Cane Road, London, W12 0NN, United Kingdom. Equal contribution (Laura E Constable, Nusrat Iqbal, Ailsa Hart, Phillip J Tozer, Nick Powell).

## Abstract

**Background and aims:** Perianal fistulization is a challenging phenotype of Crohn’s disease (CD). Unravelling the cytokine networks and cellular mediators driving tissue injury in perianal CD (pCD) will help inform much needed novel treatment strategies.

**Methods:** The phenotype of different T-cell subsets, including unconventional lineages, such as γδ T-cells, MAIT and iNKT-cells in fistula tract tissue and blood samples of patients with pCD or cryptoglandular perianal fistulas was determined using multiparameter flow cytometry. Transcriptomic profiling of fistula tract tissue was performed by RNA-sequencing.

**Results:** CD161^+^ CD4^+^ T-cells and CD161^+^ CD4^-^ CD8^-^ iNKT-cells significantly accumulated in fistula tissue and produced interleukin-(IL)22 and IL13. Transcripts associated with epithelial-to-mesenchymal transition (EMT), extracellular matrix (ECM) remodelling, interferon-gamma, JAK-STAT and lymphocyte signalling were enriched in pCD, as well as inhibition of pathways associated with wound closure. IL22-responsive transcripts were significantly enriched in fistulas and network analysis identified IL22-mediated regulation of EMT, ECM and other inflammatory pathways.

**Conclusion:** This study provides novel molecular and cellular insights into fistula pathogenesis, identifying IL22 producing lymphocytes as novel immune regulators of EMT and ECM dysregulation in perianal fistulas. Targeting the regulatory networks controlling IL22 producing lymphocytes may offer novel therapeutic strategies in pCD.

## Introduction

Crohn’s disease (CD) is a chronic immune mediated disease characterised by unpredictable, relapsing and remitting inflammation affecting any part of the gastrointestinal tract. Perianal fistulas are a common and devastating complication of CD affecting approximately 25% of patients, presenting with debilitating symptoms including perianal pain and discharge, negative impact on patient wellbeing, sleep disturbance, sexual dysfunction, and poor personal hygiene^1,2^. Perianal fistulas are challenging to treat and whilst surgical management can result in healing in >60% of patients, these procedures can carry continence risk and are often only indicated in the absence of proctitis^3^. Even with the introduction of anti-cytokine (including anti-TNF, anti-IL23), anti-integrin and cell-based therapies, a substantial proportion of patients will fail to achieve long-term fistula closure^4,5^. The development of new treatment strategies is likely to be aided by research focused on pinpointing the molecules, cells and immune networks responsible for driving tissue injury in perianal CD (pCD).

Current understanding of pCD immunopathology implicates a complex interplay between innate and adaptive immune cells, dysregulation of inflammatory and tissue repair functions, and involvement of different cytokine responses, including IL13, TNFα and IL22^6–8^. However, it is not clear how dysregulated immune responses control key biological pathways implicated in fistula pathogenesis. In the perianal fistula microenvironment, epithelial-to-mesenchymal transition (EMT) is a dominant pathway. Intestinal epithelial cells lose their cell polarity, downregulating expression of cell-cell adhesion molecules (e.g., *CDH1*)^9^, gaining migratory and invasive properties and adopt a mesenchymal cell phenotype, orchestrated by several transcriptional regulators of EMT (e.g., *SNAI1*, *ITGB6*, *ETS1*)^10,11^. Activation of myofibroblasts and induction of matrix metalloproteinases (MMPs) like MMP13, MMP9 and MMP3, coupled with a relative lack of tissue inhibitors of matrix metalloproteinases (TIMPs)^11–13^, allows cells to spread into adjacent tissues and extend into the fistula tract. *In vitro* treatment of epithelial cell lines or colonic lamina propria fibroblasts isolated from patients with pCD with TNFα or IL13, have been shown to induce expression of EMT-associated transcription factors^6–8^.

The phenotype, function and regulation of T-cells in the fistula microenvironment is of particular interest, since these potent cytokine producing cells are master regulators of tissue immune responses. Memory CD4^+^ and CD8^+^ T-cells^8,9^, including cells resembling Th1, Th17 and Th1-Th17 hybrid cells infiltrate the fistula tract wall^14^. However, it is now appreciated that T-cell biology is complex, and beyond classical T-helper lineages, unconventional lineages, including γδ T-cells, mucosal-associated invariant T-cells (MAIT) and invariant Natural Killer-like T-cells (iNKT-cells) also play important roles in the regulation of tissue immune responses. The relative contribution of the different T-cells subsets and lineages, and how these cells regulate key biological functions in the fistula microenvironment is poorly understood. This study set out to probe the phenotype and characteristics of the T-cell landscape in pCD and provide new insights into the molecular pathways they potentially regulate, to support the development of novel mechanistically informed, targeted therapeutics in the management of fistulizing CD.

## Methods

### Sample collection

Ethical approval for this study was obtained from South West-Frenchay Research Ethics Committee (REC: 19/SW/0024, IRAS Project ID: 252059). Curettage of the fistula tract was performed alongside collection of paired peripheral blood samples, from patients with CD (n=14) or cryptoglandular (n=17) perianal fistulas (**Supplementary Table.1**). Patients were classed as having cryptoglandular disease if there was no established diagnosis of CD, however patients with a presumed cryptoglandular fistula and any suspicion of IBD (erratic or frequent bowel habit, family history of IBD, any systemic or GI symptoms consistent with IBD) would go on to have investigations such as faecal calprotectin, video capsule enteroscopy, colonoscopy and/or fistula biopsy. Single biopsies were collected from the fistula tract and surrounding macroscopically uninflamed rectal mucosa (confirmed by rectal speculum examination by colorectal surgeon at the time of surgery) from patients with CD (Fistula n=11, rectal n=9) or cryptoglandular perianal fistulas (Fistula n=11, n=9) (**Supplementary Table.2**). Specimens were collected with informed consent from patients and the study was conducted in compliance with the relevant ethic regulations related to human research.

### Isolation of unfractionated cells from fistula and peripheral blood samples

Unfractionated fistula cells were isolated by enzymatic digestion. Briefly, fistula curettage specimens were incubated at 37°C in complete RPMI (Gibco) supplemented with 0.5mg/ml collagenase D, 10µg/ml DNase and 1.5mg/ml Dispase II (all Roche). Single-cell suspension was filtered through a 70μm filter to remove remaining undigested tissue. A red blood cell lysis (Biolegend) step was performed to remove erythrocytes. Cell viability was assessed using 0.4% Trypan Blue solution (Gibco). Peripheral blood mononuclear cells (PBMCs) were isolated by density-gradient centrifugation using Lymphoprep (STEMCELL Technologies). Cell viability was assessed using 0.4% Trypan Blue solution (Gibco). Isolated PBMCs were cryopreserved using fetal bovine serum (FBS; Gibco) supplemented with 10% dimethyl sulfoxide (DMSO; Sigma-Aldrich).

### Flow cytometry

Isolated PBMCs and unfractionated fistula cells were stained for surface expression of different T-cell receptor lineages TCR Vα7.2-PE-TexRed (clone 3C10), TCR γδ- APC (clone B1) and TCR Vα24-Jα18-BV510 (clone 6B11) (all Biolegend). Cells were washed with PBS (Gibco) and centrifugation at 1800rpm, 4°C for 5min prior to stimulation in complete RPMI-1640 supplemented with 50ng/ml phorbol 12-myristate 12-acetate (PMA; Sigma-Aldrich), 1µg/ml ionomycin (Sigma-Aldrich), 2µM monensin (Invitrogen) and 5µg/ml Brefeldin A (Invitrogen). PBMCs were stimulated for 5hr and unfractionated fistula cells for 3hr. Stimulated cells were stained for cell surface expression of CD45-BV650 (Biolegend, clone HI30), CD3-FITC (Biolegend, clone OKT3), CD4-APC-CY7 (Biolegend, clone OKT4), CD8-BUV395 (BD Biosciences, clone RPA-T8), CD161-BV786 (Biolegend, clone HP-3G10), CD56-PE-Cy5 (Biolegend, clone 5.1H11) and fixable viability stain-AF700 (BD Biosciences). Cells were fixed and permeabilized using the Foxp3 fixation/permeabilization buffer kit (eBioscience), and intracellular cytokine staining (ICS) performed for IFNγ-BUV737 (BD Biosciences, clone 4S.B3), Granzyme B-PE (Biolegend, clone QA16A02), IL17A- BV421 (Biolegend, clone BL168), IL13-BV711 (BD Biosciences, clone JES10-5A2), IL22-PE-Cy7 (Biolegend, clone 2G12A41) and TNFα-BV605 (Biolegend, clone MAb11). Cells were resuspended in PBS and stored overnight in the dark at 4°C until acquisition. Data were acquired on a BD LSRFortessa flow cytometer, recorded using BD FACSDiva 6.0 (BD Biosciences) and analysed using FlowJo (Treestar Inc).

### RNA extraction and library preparation

Biopsies were collected into RNAlater (Invitrogen) and stored for 24hr at 4°C before long-term storage at -80°C. Biopsies were homogenized and lyzed with Qiazol (QIAGEN) and RNA extracted using chloroform (Sigma-Aldrich). RNA was precipitated with isopropanol (Sigma-Aldrich) and GlycoBlue (Gibco) before being resuspended in RNase-free water (Gibco). RNA quality and quantity was assessed by Nanodrop, Qubit 3.0 fluorometer (Life Technologies) and 2100 Bioanalyzer (Agilent). RNA was stored at -80°C. Paired-end libraries were generated using the NEBNext^®^ Ultra™ RNA Library Prep Kit for Illumina^®^, then sequenced on the HiSeq Illumina platform (Novogene) and deposited on NCBI GEO (series GSE228391).

### RNA-seq data analysis

Adaptor contamination and poor-quality bases were removed using trimmomatic (v. 0.39)^15^. The resulting reads were mapped to the human genome (GRCh38) using Hisat2 (v. 2.2.1) with default parameters^16^. The number of reads mapping to the genomic features annotated in Ensembl^17^ with MAPQ ≥ 10 was calculated for all samples using htseq-count (v. 0.11.3)^18^ with default parameters. Data for Ensembl genes with no associated ENTREZ gene identifier were discarded; counts for Ensembl genes mapped to the same ENTREZ gene identifier were summed up samplewise.

Differential expression analysis was performed in R (v. 3.6.1)^19^ using DESeq2^20^ and adjusting for (i) the date of RNA extraction and the sequencing batch in the comparison between fistula samples from Crohn’s disease patients and cryptoglandular donors; or for (ii) donor identity in the comparison between fistula and rectal samples. For each contrast, only genes with at least 3 counts per million in at least 5 samples were tested. The Benjamini and Hochberg procedure^21^ was applied for multiple testing correction.

Enrichment of signatures in MSigDB^22,23^ was assessed using GSEA^24^. Genes were ranked by decreasing scores calculated by taking the geometric mean between the absolute value of the log fold change and the p-value from DEseq2 following log10 transformation and change of sign. The sign of the log fold change was finally multiplied to the ranking measure. These calculations were executed in R (v. 3.6.1) using the packages msigdbr and fsgsea with default parameters. Cytokine-responsive gene lists were downloaded from previous studies^25,26^ and their enrichment in individual transcriptome profiles evaluated with the GSVA^27^ package using the ssGSEA method without score scaling. VIPER^28^ was used to infer transcription factor activities based on the interactions DoRothEA^29^ reports with confidence encoded as A, B or C. Genes were ranked based on the p-values and sign of the test statistics from DESeq2. Enrichment was calculated for regulons with at least 5 target genes and adjusted for pleiotropic effects. The results of differential gene expression analysis comparing fistula and rectal samples in CD were analysed with ViralLink^30^. The resulting network was filtered to Include only members of the IL22 pathway listed in BioCarta^31^, and downstream transcription factors and differentially expressed genes (DEGs) identified in the analyses described above. DEGs with FDR >0.1 and transcription regulators with no connection in the network were discarded. Network visualizations were created in Cytoscape^32^.

### Statistical analysis

Data are expressed as medians or mean ± SD where specified. Hypothesis testing was performed using paired Wilcoxon signed rank tests or unpaired t-tests, where appropriate, using GraphPad Prism 10.0 (GraphPad Inc., USA).

## Results

### Conventional CD4^+^ T-cells and invariant natural killer T-cells are expanded in perianal fistulas

To investigate the T-cell and molecular landscape of pCD we performed multiparameter flow cytometry of cells isolated from fistula tract curettage, and peripheral blood, and performed next generation RNA-sequencing of biopsies sampled from the fistula tract of patients with pCD and cryptoglandular fistulas (**Figure 1A**). Profiling the phenotype of CD3^+^ T-cells from patients with perianal fistulizing CD (n=14) and cryptoglandular perianal fistulas (n=17) we demonstrated the presence of five main T-cell subsets, including conventional CD4^+^ and CD8^+^ T-cells, iNKT-cells, MAIT-cells and γδ T-cells (**Figure 1B**). t-SNE plots based on the median intensity of cell surface marker expression in blood and fistula samples showed that CD4^+^ and CD8^+^ T-cells comprised the majority of CD3^+^ T-cells (**Figure 1C**). Analysis of the proportional abundance of the different subsets demonstrated that CD4^+^ T-cells and iNKT-cells were significantly increased in fistula tissue compared to peripheral blood. By contrast, the proportion of CD8^+^ and γδ T-cells was reduced in fistulas in comparison to peripheral blood, whilst the abundance of MAIT-cells was unchanged. The overall proportional abundance of different T-cell subsets was similar in pCD and cryptoglandular fistulas (**Figure 1D**), although the most consistent finding was relative enrichment of iNKT-cells and CD4^+^ T-cells in both CD and cryptoglandular fistulas (**Figure 1E**).

**Figure 1:**
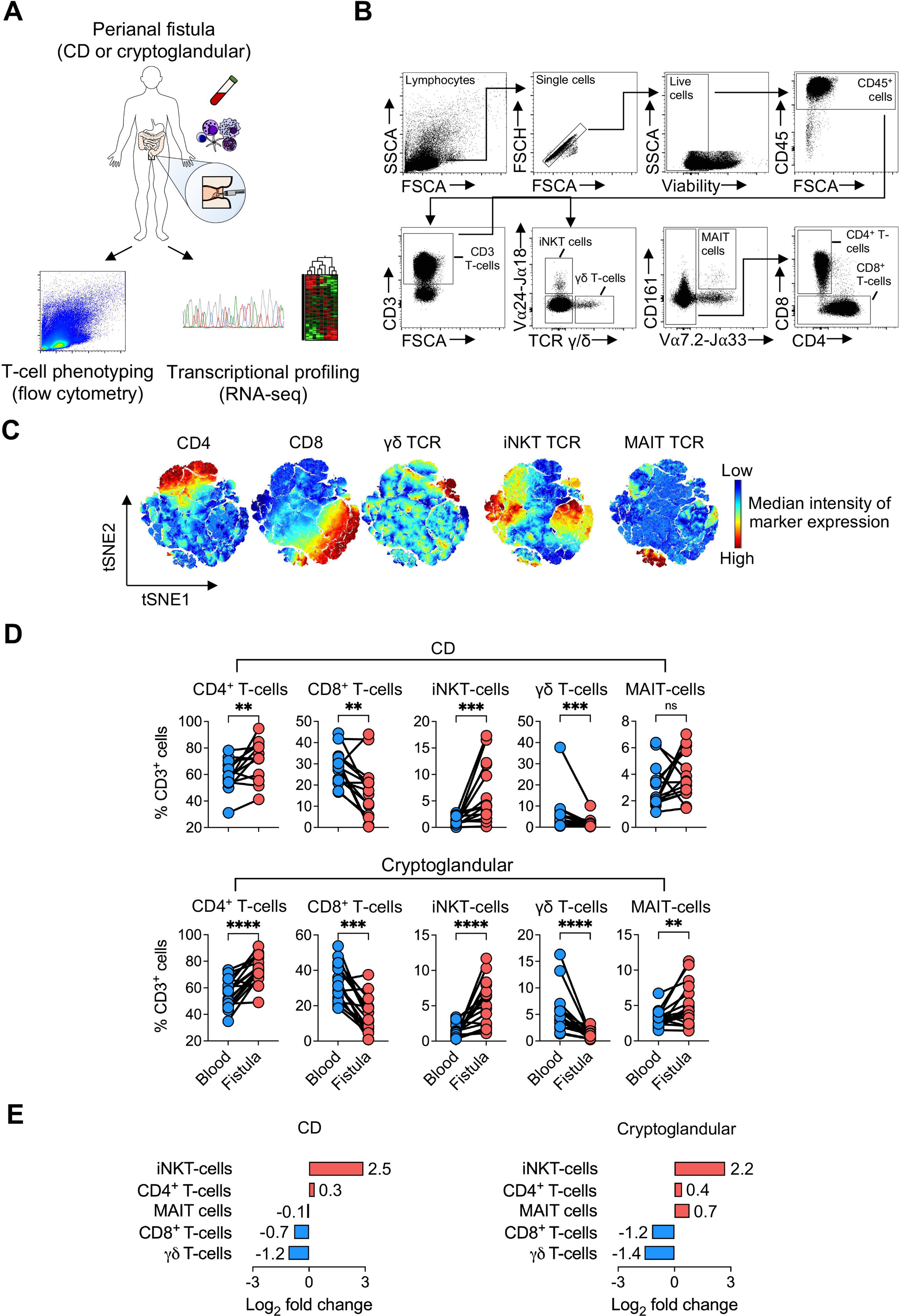
CD4^+^ T-cells and invariant natural killer T-cells are expanded in perianal fistulas. A) Schematic representation of multiparameter flow cytometry experiments phenotyping CD3^+^ T-cells in paired peripheral blood and fistula samples (pCD=14, cryptoglandular n=17) and RNA-sequencing of fistula (n=13) and rectal (n=10) biopsies collected from patients with pCD or cryptoglandular fistulizing disease. B) Flow cytometry gating strategy used to identify conventional and unconventional T-cell subsets in peripheral blood and fistula tissue. C) tSNE plot of CD3^+^ T-cells (gated on live CD45^+^ cells) showing expression of CD4, CD8, γδ TCR, iNKT TCR and MAIT TCR. D) Quantification of T-cell subsets in peripheral blood and fistula tissue in pCD (upper panel) or cryptoglandular (lower panel) fistulizing disease. E) Log2 fold change in the proportion of T-cell subsets in pCD and cryptoglandular fistula compared to peripheral blood. Statistical significance was calculated using paired Wilcoxon signed rank test. **P<0.01, ***P<0.001, ****P<0.0001, ^ns^non-significant.

### Increased frequency of IL22 producing CD4^+^ T-cells and iNKT-cells in perianal fistulas

Next, we examined production of key cytokines, including IFNγ, IL13, IL17A, IL22, TNFα and granzyme B by the T-cell compartment in the fistula microenvironment. Overall, in comparison with peripheral blood, the most striking difference in fistula tissue was a significant increase in the proportional abundance of CD3^+^ T-cells expressing IL22 and IL13 (**Figure 2A and 2B**). Similar enrichment of these responses was observed in both CD and cryptoglandular fistulas (**Figure 2C and 2D**). The chief producers of IL22 and IL13 in fistula tissue were conventional CD4^+^ T-cells and iNKT-cells (**Figure 2E**).

**Figure 2:**
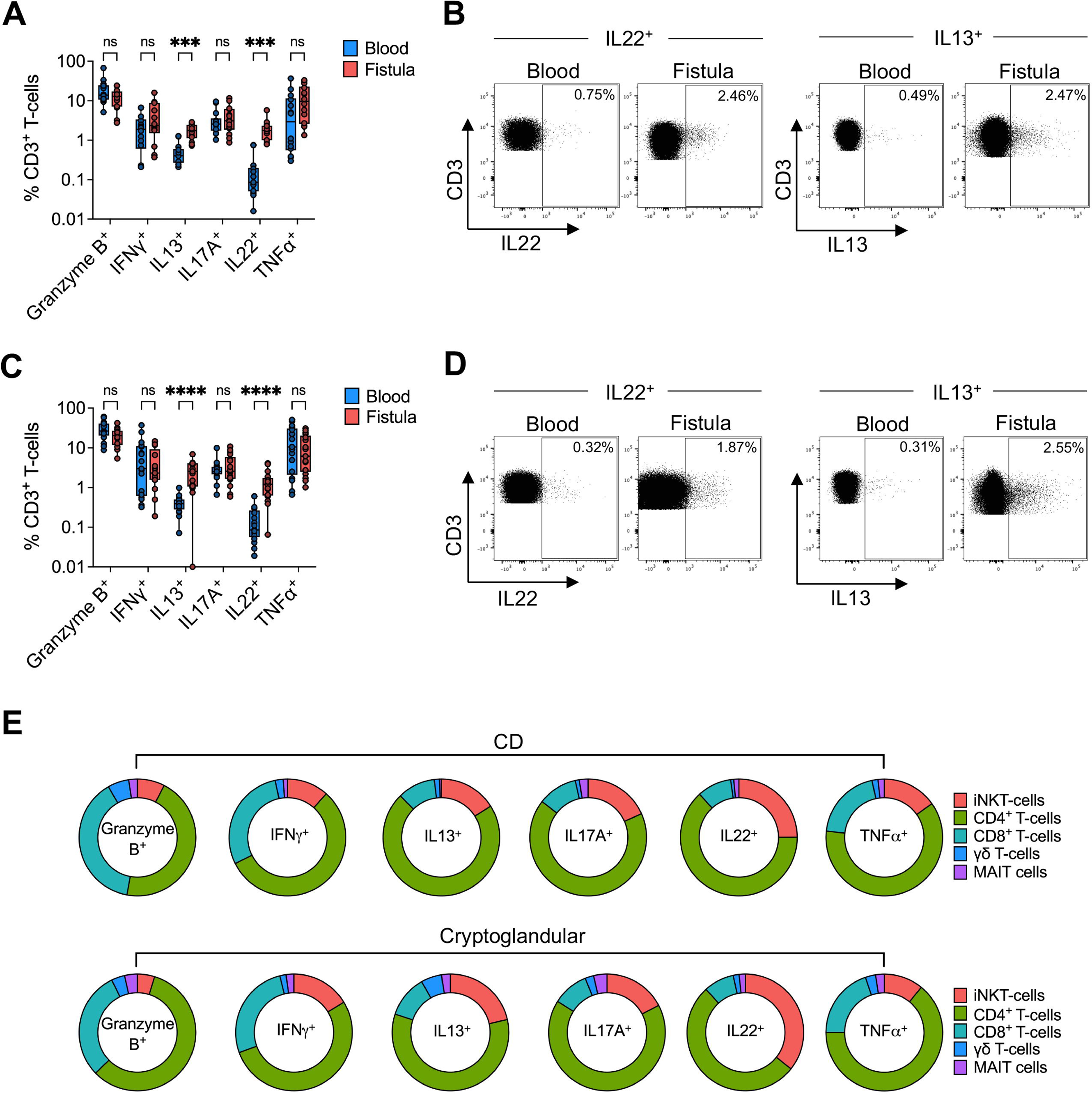
CD4^+^ T-cells and iNKT-cells are a major cellular source of IL22 and IL13 in fistula tissue. A) Frequency of granzyme B, IFNγ, IL13, IL17A, IL22 and TNF⍺ producing CD3^+^ T-cells in paired peripheral blood and fistula tissue from patients with pCD (n=14). B) Representative flow cytometry plot showing IL22 and IL13 production by CD3^+^ T-cells in blood and fistula samples from pCD patients. C) Frequency of granzyme B, IFNγ, IL13, IL17A, IL22 and TNF⍺ producing CD3^+^ T-cells in paired peripheral blood and fistula tissue from patients with cryptoglandular fistulizing disease (n=17). D) Representative flow cytometry plot showing IL22 and IL13 production by CD3^+^ T-cells in blood and fistula samples from patients with cryptoglandular fistulizing disease. E) Frequency of iNKT-cells, CD4^+^ T-cells, CD8^+^ T-cells, γδ T-cells and MAIT cells as a proportion of CD3^+^ granzyme B^+^, IFNγ^+^, IL13^+^, IL17A^+^, IL22^+^ or TNF⍺^+^ T- cells in pCD and cryptoglandular fistula tissue. Data shown as medians. Statistical significance calculated using paired Wilcoxon signed rank test. \*\*\*\**P*<0.0001, ^ns^non- significant.

Similarly, at a cell population level, there was a significant increase in the proportion of IL22^+^ and IL13^+^ producing CD4^+^ T-cells in both CD and cryptoglandular fistulas (**Figure 3A, 3B, Supplementary Figure 1A and 1B**). iNKT-cells are known to exist as functionally distinct subsets based on their expression of CD4 and CD8 co-receptors (ref). In fistula tissue, double negative (DN; CD8^-^ CD4^-^) iNKT-cells were significantly increased, and CD4^+^ and CD8^+^ subsets reciprocally decreased in frequency in both CD and cryptoglandular fistulas (**Figure 3C, 3D**, **Supplementary Figure 1C and 1D**). A significant increase in the proportion of IL22^+^ producing DN iNKT-cells was also seen in CD and cryptoglandular fistulas (**Figure 3E, 3F, Supplementary Figure 1E and 1F**).

**Figure 3:**
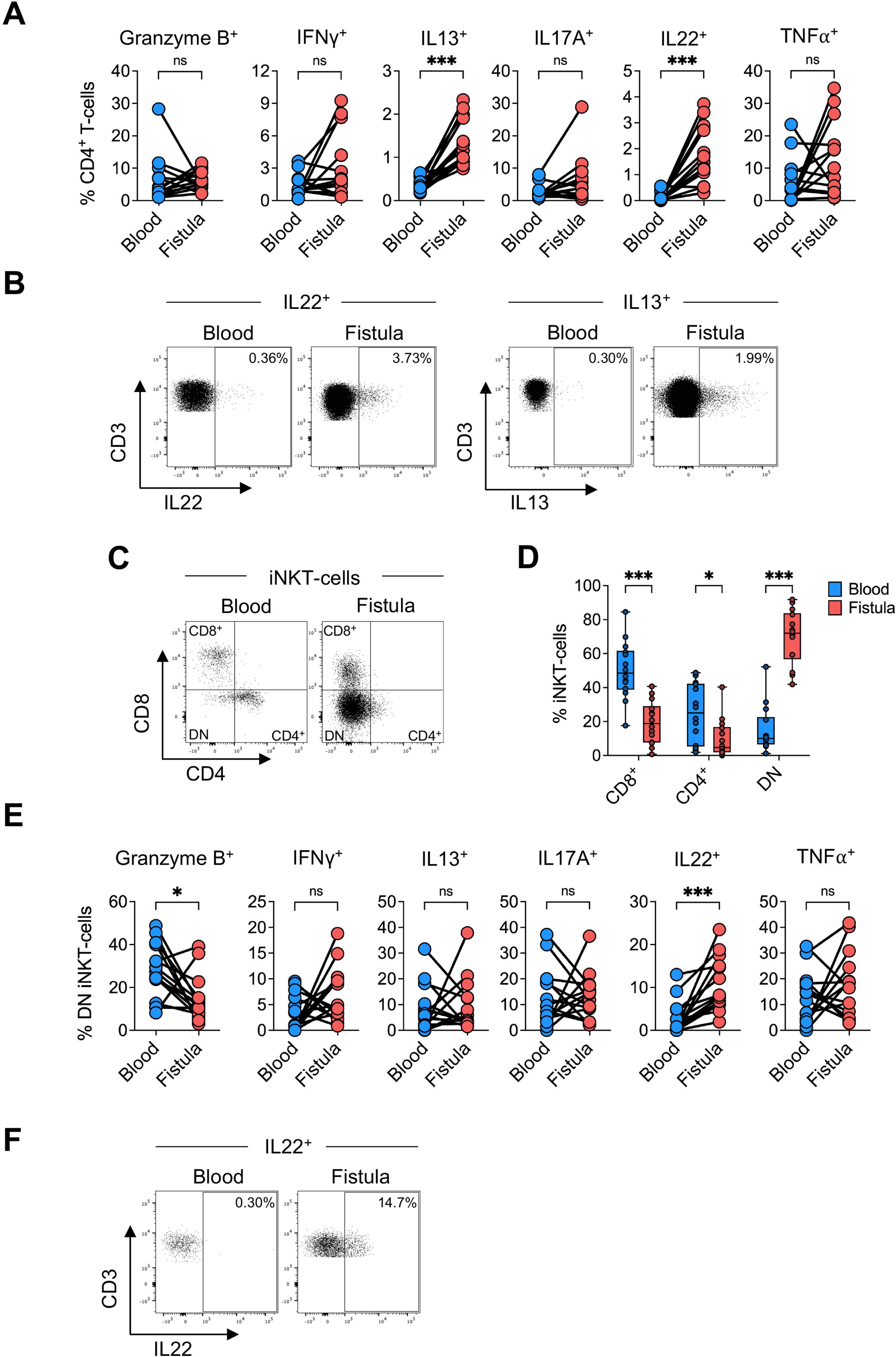
IL22 and IL13 producing CD4^+^ T-cells and IL22 producing iNKT-cells are increased in fistula tissue in pCD. A) Proportion of cytokine-producing CD4^+^ T-cells in paired peripheral blood and fistula tissue in pCD patients (n=14). B) Representative flow cytometry plot showing production of IL22 and IL13 by CD4^+^ T-cells in blood and fistula tissue. C) Representative flow cytometry plot showing the division of iNKT-cells into CD8^+^, CD4^+^ and CD8^-^ CD4^-^ (DN) subsets. D) Frequency of each iNKT-cell subset in blood and fistula tissue in pCD patients. E) Proportion of cytokine producing DN iNKT-cells in blood and fistula samples in pCD patients. F) Representative flow cytometry plots showing production of IL22 by DN iNKT-cells in blood and fistula. Data expressed as medians. Statistical significance calculated using paired Wilcoxon signed rank test. \**P*<0.05, \*\*\**P*<0.001, ^ns^not statistically significant.

CD161 marks T-cells with superior capacity for proinflammatory cytokine production in IBD^33^. The proportion of cells expressing CD161 was enriched in fistula tissue in comparison with blood across most of the T-cell subsets, except MAIT cells (**Supplementary Figure 2A and 2B**). In fistula tissue, there was a significantly increased proportion of cells producing IL17A, IL22 and TNFα by the CD161^+^ subset of CD4^+^ T-cells in comparison with their CD161^-^ counterparts (**Supplementary Figure 2C and 2D**).

Similarly, there was a significantly increased fraction of IFNγ, IL13, IL17A and TNFα producing iNKT-cells in the CD161^+^ compartment in comparison with CD161^-^ cells. A trend towards increased IL22 production by CD161^+^ iNKT-cells was also observed (p=0.06) (**Supplementary Figure 2E and 2F**).

The frequency of CD161^+^ CD4^+^ T-cells producing IL13 and IL22 and CD161^+^ DN iNKTs producing IL22 were proportionally expanded in fistula tissue compared to peripheral blood (**Supplementary Figure 2G and 2H**). Similar findings were observed in cryptoglandular fistulas, with relative expansion of IL22 producing CD161^+^ CD4^+^ T-cells and DN iNKT-cells (**Supplementary Figure 3**).

### The transcriptional architecture of perianal fistulas is distinct to rectal mucosa and characterised by epithelial-to-mesenchymal transition, dysregulation of the extracellular matrix and immune activation

To identify molecular mechanisms characterising the perianal fistula environment, we analysed gene expression profiles in biopsies from patients with pCD (fistula n=11, rectal n=9) or cryptoglandular (fistula n=11, rectal n=9) fistulizing disease. Principal component analysis (PCA) showed a strong correlation between the first principal component and sample tissue of origin (Spearman’s rho=0.79) (**Figure 4A**), indicating that tissue origin was the largest source of variation. Hierarchical clustering of differentially expressed genes (DEGs) (FDR<0.05) between fistula and rectal tissue in both diseases further underscored the pronounced transcriptional changes between tissues, but strikingly highlighted remarkable similarities in the transcriptome of pCD and cryptoglandular fistulas (**Figure 4B**). Many genes were differentially expressed between fistula and rectal tissue in both conditions (**Figure 4C**). The most significantly and highly upregulated DEGs in pCD and cryptoglandular fistulas encoded proteases (*MMP13, ADAMTS18, KLK7)*, protease inhibitors (*SERPINB13, SERPINB3, SERPINB4*), extracellular matrix proteins (*COMP, COL11A1, ACAN)*, Wnt signalling molecules (*WNT2, SFRP4, ISM1*), transcriptional regulators of tissue morphogenesis (*HOXC10, HOXC11, HOXC6*) and antimicrobial responses (*LBP, S100A7, PGLYRP4)*. Significantly downregulated genes included those encoding epithelial cell structure and function (e.g., *CLDN8, CDHR5, MUC17*), lipid metabolism (*AQP8, FABP1, HNF4A*) and transmembrane ion transport (*BEST2, CLCA1, SLC26A3*).

**Figure 4:**
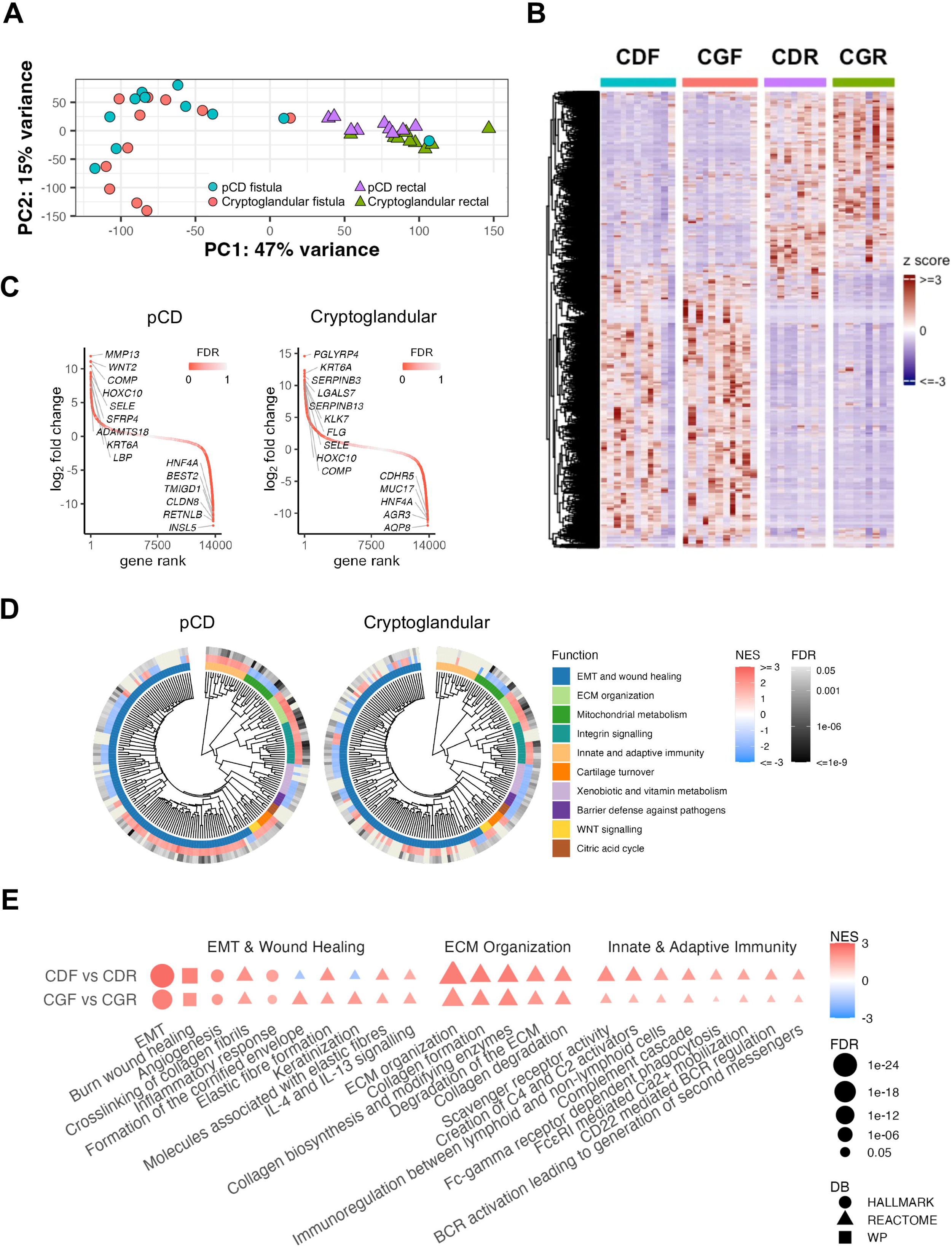
Fistulas are characterised by activation of epithelial-to-mesenchymal transition and organization of the extracellular matrix. A) PCA plot of fistula (n=22) and rectal samples (n=18) from patients with pCD or cryptoglandular (CG) disease. B) Heatmap of z-score transformed transcripts per millions for genes differentially expressed (FDR < 0.05) comparing fistula vs rectal samples in pCD or CG disease. C) Distributions of gene expression changes between fistula and rectal tissue in pCD and CG disease. D) Hierarchical clustering of pathways regulated in significantly different ways (FDR < 0.05) in fistula versus rectal tissue in pCD and CG disease.Circular dendrogram shows pathway similarities quantified by Jaccard scores of their gene lists. Inner, middle and outer rings represent functional categories, normalized enrichment scores and FDR from GSEA, respectively. E) Pathways involved in epithelial-to-mesenchymal transition and wound healing, ECM organization and innate and adaptive immunity, which exhibit significantly altered activity (FDR <0.05) in fistula versus rectal tissue in pCD or CG disease.

Gene set enrichment analysis (GSEA) of gene expression changes between fistula and rectal tissue in pCD and cryptoglandular fistulizing disease identified 211 biological pathways as significantly perturbed (FDR <0.05) across the two aetiologies. These could be grouped into ten functional categories (**Figure 4D and Supplementary Table 3**). The largest cluster included pathways involved in EMT and wound healing, as well as other related processes like angiogenesis, formation of elastic fibres and the inflammatory response, particularly interleukin-4 and interleukin- 13 signalling, and were significantly more activated in fistula than rectal tissue (**Figure 4E**). Although activated in both pCD and cryptoglandular fistulas, EMT, inflammatory response and wound healing pathways were at least two orders of magnitude more significantly impacted in CD than cryptoglandular disease. Complementary analyses further emphasized the relevance of EMT in pCD, showing reciprocal increase in activity of transcription factors associated with induction of EMT (*ETS2, ERG, SMAD3*) and reduction in those associated with the maintenance of an epithelial cell phenotype (*ZEB2, ELF3, ELF5, EHF*) (**Supplementary Figure 4A**). Except for *ERG*, similar enrichment of EMT-associated transcription factors was observed comparing cryptoglandular fistula to rectal tissue. However, other transcription factors regulating lymphocyte activation and cytokine signalling (*SPI1, LYL1, IKZF1, STAT1)* were more active in pCD than cryptoglandular fistula tissue. *AHR*, a transcriptional regulator of IL22 expression, was equally active in pCD and cryptoglandular fistulas compared to rectal tissue. Furthermore, many activated transcription factors, especially those involved in lymphocyte activation and cytokine signalling, form a gene regulatory network converging on *ERG* (**Supplementary Figure 4B**). Other key cellular functions impacted in fistulas included remodelling and organization of the ECM, particularly collagen formation and degradation (**Figure 4E**). Pathways like keratinization and formation of the cornified envelope, involved in wound re-epithelization and closure, were significantly more active in fistula compared to rectal tissue in cryptoglandular, but not fistulizing pCD (**Figure 4E**). Furthermore, innate immune pathways like complement cascade, macrophage-mediated phagocytosis and B-cell activation were significantly activated (FDR<0.05) in pCD but not cryptoglandular fistulas.

### Expression of matrisome related genes and proteases indicate ECM remodelling is a key feature of both CD and cryptoglandular fistulas

To further explore the role of the ECM and its remodelling in fistulizing disease in greater detail, we also investigated the transcriptional phenotype of perianal fistulas by focusing on genes associated with different components of the ECM termed the “matrisome”^34^. In both pCD and cryptoglandular fistulas, genes coding for basement membrane proteins, collagens, glycoproteins, and proteoglycans were expressed at significantly higher levels in fistula compared to the rectal tissue (**Figure 5A**). Additionally, genes encoding ECM regulators, particularly enzymes and proteases involved in tissue remodelling and secreted factors that bind to the ECM, also exhibited the same phenotype. Only ECM-affiliated genes, those sharing architectural similarities or known to be associated with ECM proteins, were not found to be preferentially expressed in fistula tissue.

**Figure 5:**
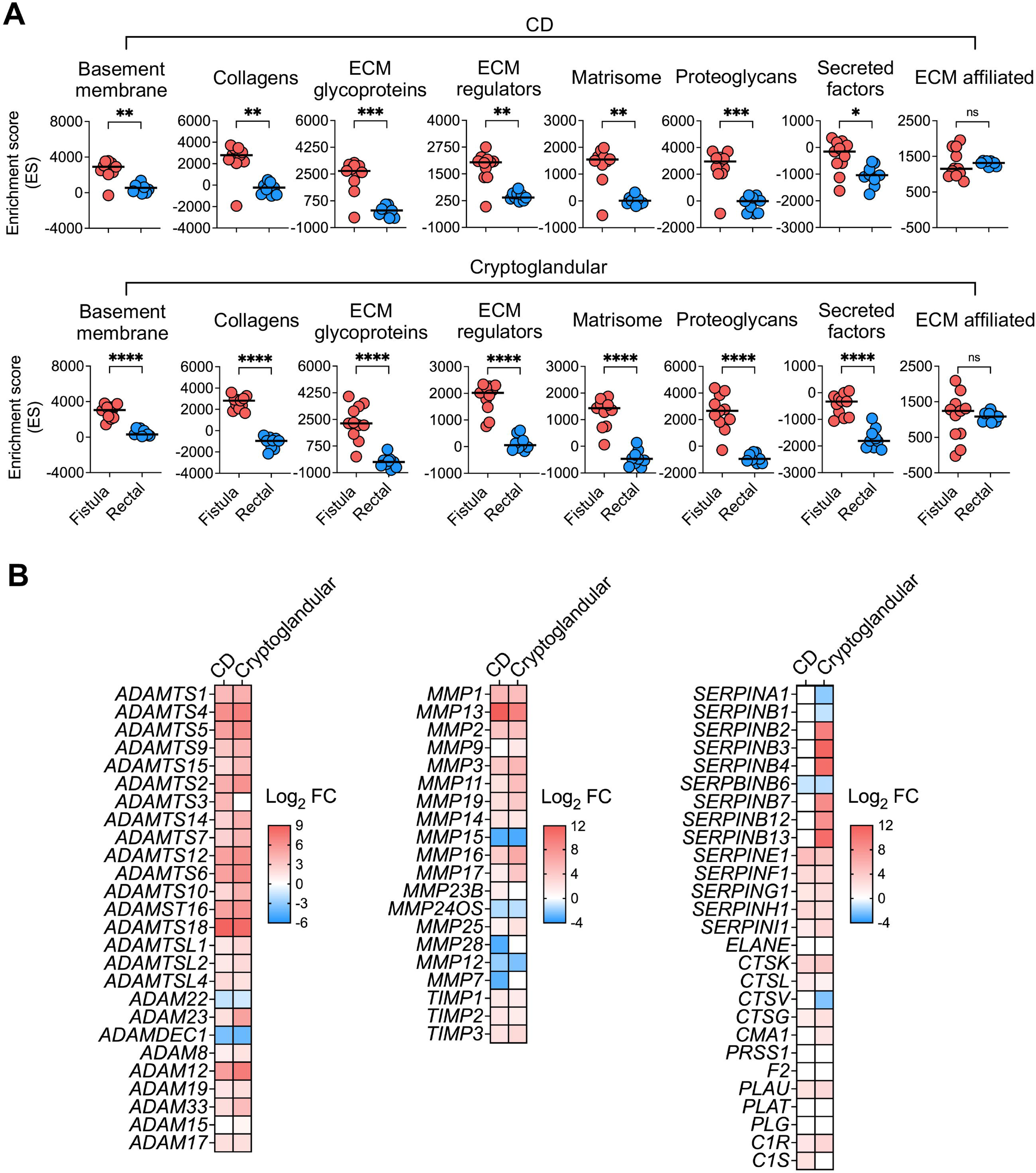
Dysregulated ECM responses and tissue remodelling is a key feature of perianal fistulas. A) Enrichment score distributions of gene signatures encoding constituents of the ECM in fistula and rectal samples estimated by single sample GSEA. B) Differences in gene expression levels for genes encoding metalloproteinases and their inhibitors based on the comparisons of pCDF vs pCDR samples and CGF vs CGR samples. Data plotted for metalloproteinases of the ADAM, ADAMTS and MMP families, as well as of members of the TIMP and SERPIN families of inhibitors. Statistical significance of the differences in median values performed using Mann-Whitney tests; \**P*<0.05, \*\**P*<0.01, \*\*\**P*<0.001, \*\*\*\**P*<0.0001, ^ns^non- significant.

Given the expression of ECM regulators were preferentially enriched in fistulas, we investigated the change in expression of several classes of metalloproteinases, endopeptidases and their inhibitors in fistula and rectal tissue in pCD and cryptoglandular fistulizing disease. We found most matrix metalloproteinases (MMPs) to be equally highly over-expressed in pCD and cryptoglandular fistulas compared to rectum (**Figure 5B**). However, unlike other studies^12^, we demonstrated a simultaneous overexpression of tissue inhibitors of matrix metalloproteinases (TIMPs) in both pCD and cryptoglandular fistulas, being 3- to 4-fold upregulated in fistula tissue. Similarly, other proteases of the ADAM/ADAMTS family involved in ECM turnover demonstrated similar expression patterns in pCD and cryptoglandular fistulas, with the majority being upregulated in fistula compared to rectal tissue. The expression of serine proteases and their natural inhibitors (SERPINs) however, did appear to be regulated differently in pCD and cryptoglandular disease. Whilst many serine proteases were expressed at similar levels between CD and cryptoglandular fistulas, including members of the cathepsin family (*CTSK, CTSL, CTSG*), expression of their natural inhibitors in the SERPIN B family could only be detected in cryptoglandular fistula tissue.

### Enhanced cytokine signalling and impaired wound healing pathways differentiate CD from cryptoglandular fistulas

Few studies have compared the immunopathology of pCD and cryptoglandular fistulizing disease. Our results comparing fistula to rectal tissue revealed significant overlap in their molecular pathology, particularly in relation to the ECM and tissue remodelling responses (**Figure 4D and 4E**). A genome-wide comparison between fistula and rectal tissue in pCD and cryptoglandular disease demonstrated very strong correlations between changes in both gene expression levels (Spearman’s rho≍0.89) and biological pathway activities (Spearman’s rho≍0.86) (**Figure 6A**). Furthermore, direct comparison of the transcriptome of pCD and cryptoglandular fistula tissue identified few differentially expressed (FDR<0.05) genes (**Figure 6B**). The most highly overexpressed genes included members of the TNF superfamily (*LTA*, *LTB*, *TNFSF13*), of the T- and B-cell receptor signalling and activation pathways (*IGVL5- 37*, *CD3D*, *IL2RA*, *CD40LG*) and molecules involved in cell-adhesion and T-cell trafficking (*CD52, CCR7, HLA-DPB1*).

**Figure 6:**
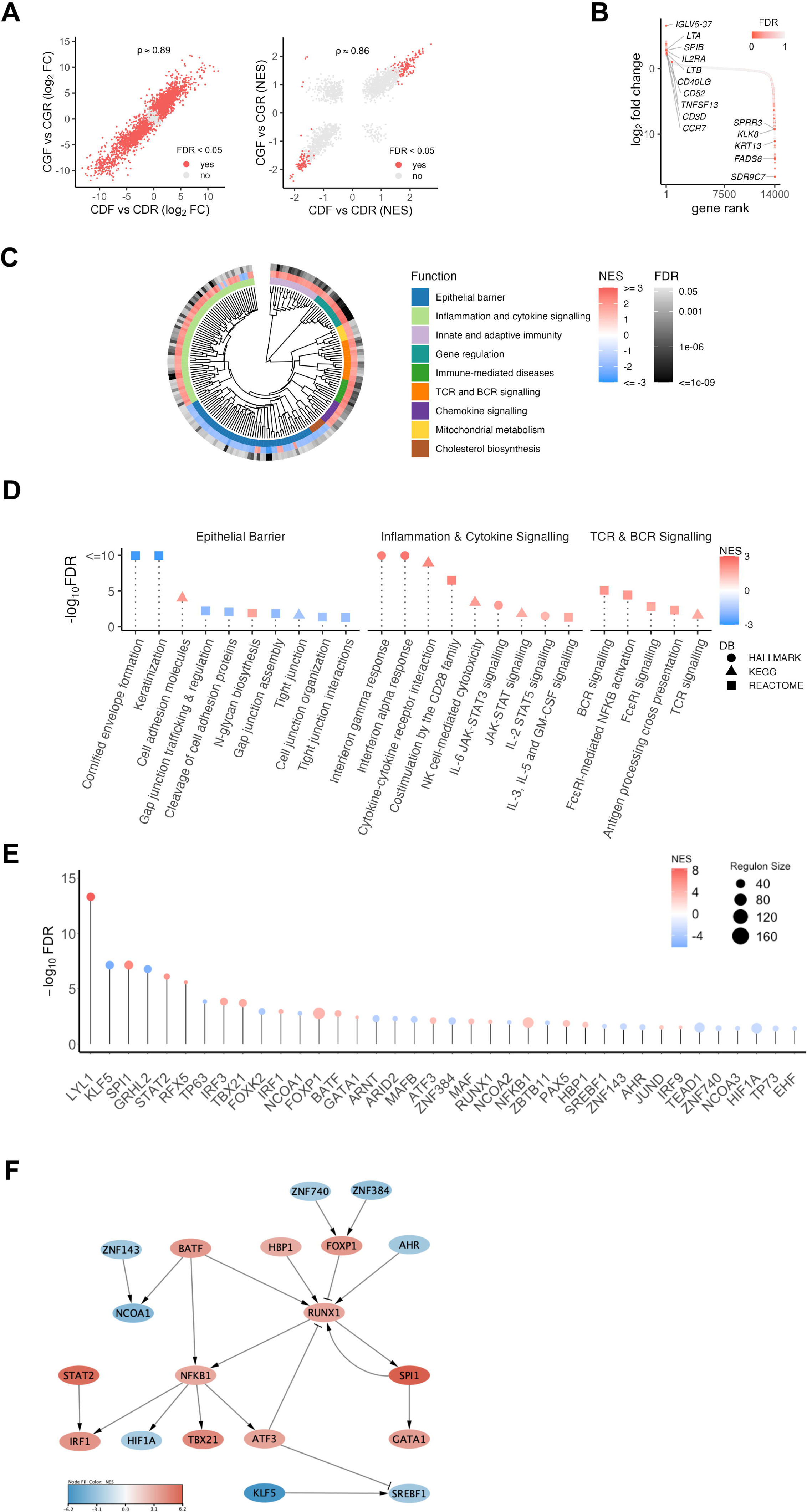
Epithelial barrier breakdown, inflammatory cytokine signalling, and B- and T-cell receptor signalling pathways are more enhanced in CD fistulas than cryptoglandular fistula. A) Relationship between gene expression changes (log2 fold changes, left panel) and biological pathway activities (normalized enrichment scores, right panel) comparing fistula vs rectal tissue in pCD (x axis) and cryptoglandular (y axis) samples. Spearman correlation coefficient shown at the top of each panel. Points are in red if the corresponding genes or pathways are significantly impacted in at least one comparison, and in light grey otherwise. B) Distribution of gene expression changes between fistula tissue in donors with pCD and cryptoglandular fistulizing disease. C) Hierarchical clustering of pathways regulated in significantly different ways (FDR < 0.05) in fistula tissue from donors with pCD and cryptoglandular fistulizing disease. Circular dendrogram shows pathway similarities quantified by Jaccard scores, while inner, middle and outer rings represent assignments to broad functional roles, normalized enrichment scores and false discovery rate, respectively. D) Pathways involved in the epithelial barrier, inflammation and cytokine signalling and B- (BCR) and T-cell receptor (TCR) signalling that exhibit altered activity in pCD versus cryptoglandular fistula tissue (FDR<0.05). E) Significant (FDR<0.05) differences in transcription regulator activities between pCD and cryptoglandular fistula tissue. Positive and negative NES values indicate increased and reduced activity in pCD fistula vs cryptoglandular fistula tissue. F) Network of regulatory interactions among genes in E); edges with triangular and T-shaped ends indicate stimulatory and inhibitory relationships, respectively.

Gene set enrichment analysis identified 189 biological pathways to be significantly impacted (FDR<0.05) comparing pCD to cryptoglandular fistula tissue. Hierarchical clustering showed that these pathways could be grouped into nine functional classes (**Figure 6C and Supplementary Table 4**). Several pathways associated with epithelial barrier structure and function including gap and tight junction assembly, trafficking, organization and interactions were significantly inhibited in pCD compared to cryptoglandular fistulas (**Figure 6D**). Highly activated pathways included those related to inflammation and cytokine signalling, particularly IFNγ and IFNα responses. Enhanced activation of JAK-STAT pathways, including IL6-STAT3 and IL2-STAT5 signalling was observed in pCD fistulas. Signalling by IL3, IL5 and GM-CSF was also significantly more activated in CD compared to cryptoglandular fistulas. Additionally, biological pathways associated with T- and B-cell receptor signalling, antigen presentation and immunoglobulin receptor signalling were also more active in pCD. Consistent with enhanced activity of interferon signalling pathways, IFNγ (*TBX21*, *IRF3*, *IRF9*) and IFNα/β (*IRF1*, *STAT2*) related transcription factors were more activated in pCD compared to cryptoglandular fistulas (**Figure 6E**). Other transcription factors associated with immune cell differentiation and activation of T-, B-cells and myeloid cells (*SPI1, GATA1, RUNX1*), were also significantly more active in pCD. Gene regulatory network analysis indicated these processes were downstream of NFκB and *RUNX1* activation (**Figure 6F**).

Unlike the comparison between fistulas, a greater magnitude of difference was observed in the transcriptional profile of rectal tissue between patients with pCD or cryptoglandular fistulizing disease. Many genes were differentially expressed and amongst the most significantly upregulated in pCD included those associated with antimicrobial responses (*REG1A, DEFA5, DEFA6*), lymphocyte activation (*CD19, CCR7, CTLA4*) and cytokine and chemokine production (*NOS2, IDO1, CCL19, CCL22*) (**Supplementary Figure 5A**). In keeping with this, pathway analysis indicated that rectal tissue in pCD was characterised by greater activation of pathways related to cytokine signalling and immune activation (**Supplementary Figure 5B and Supplementary Table 5**). Amongst the pathways most significantly activated in the rectal mucosa of pCD patients compared to cryptoglandular were the IL6-JAK-STAT3 and IL2-STAT5 signalling pathways, as well as IFNγ and IFNα responses (**Supplementary Figure 5C**). Notably, TGFβ signalling was inhibited. Several pathways associated with immune cell activation including TCR signalling, co-stimulation by CD28, PD1 signalling, and antigen processing and presentation were also more activated in rectal tissue in pCD. Preferential activation of the complement cascade and other related processes like Fcγ receptor dependent phagocytosis and activation were also observed in pCD. Furthermore, like in pCD fistulas, a greater inhibition of pathways related to cell and tight junction organization as well as keratinization were seen in CD compared to cryptoglandular rectal tissue.

### IL22 is a key transcriptional regulator of EMT and ECM remodelling in perianal fistulas

Several cytokines have been hypothesized to regulate EMT and tissue remodelling^7,8,11^. Since EMT and tissue remodelling were prominent biological mechanisms activated in pCD and cryptoglandular fistulas in this study, we next asked how they were regulated, and whether the cytokines produced by CD4^+^ T-cells and iNKTs that accumulated in fistula tissue were involved. To this end, we estimated the summarised expression level of gene signatures characterising the transcriptional phenotype of colonic epithelial organoids (“colonoids”) treated with canonical T-cell cytokines^25,26^ in our dataset. This analysis showed that IL13 and IL22 responsive genes were significantly more expressed in fistula tissue in pCD and, by contrast, IL17A responsive genes were reduced (**Figure 7A**). In cryptoglandular fistulas, in addition to upregulation of IL22 and IL13 responsive genes, significant enrichment of IFNγ and TNFα responsive genes was also observed. Furthermore, in cytokine-treated colonoids we found that multiple cytokines could impact the expression of matrisome components that we found preferentially enriched in fistula tissue. We demonstrated significant enrichment of ECM and matrisome related signatures in colonoids treated with IFNγ, IL13, IL22 and TNFα (**Figure 7B**).

**Figure 7:**
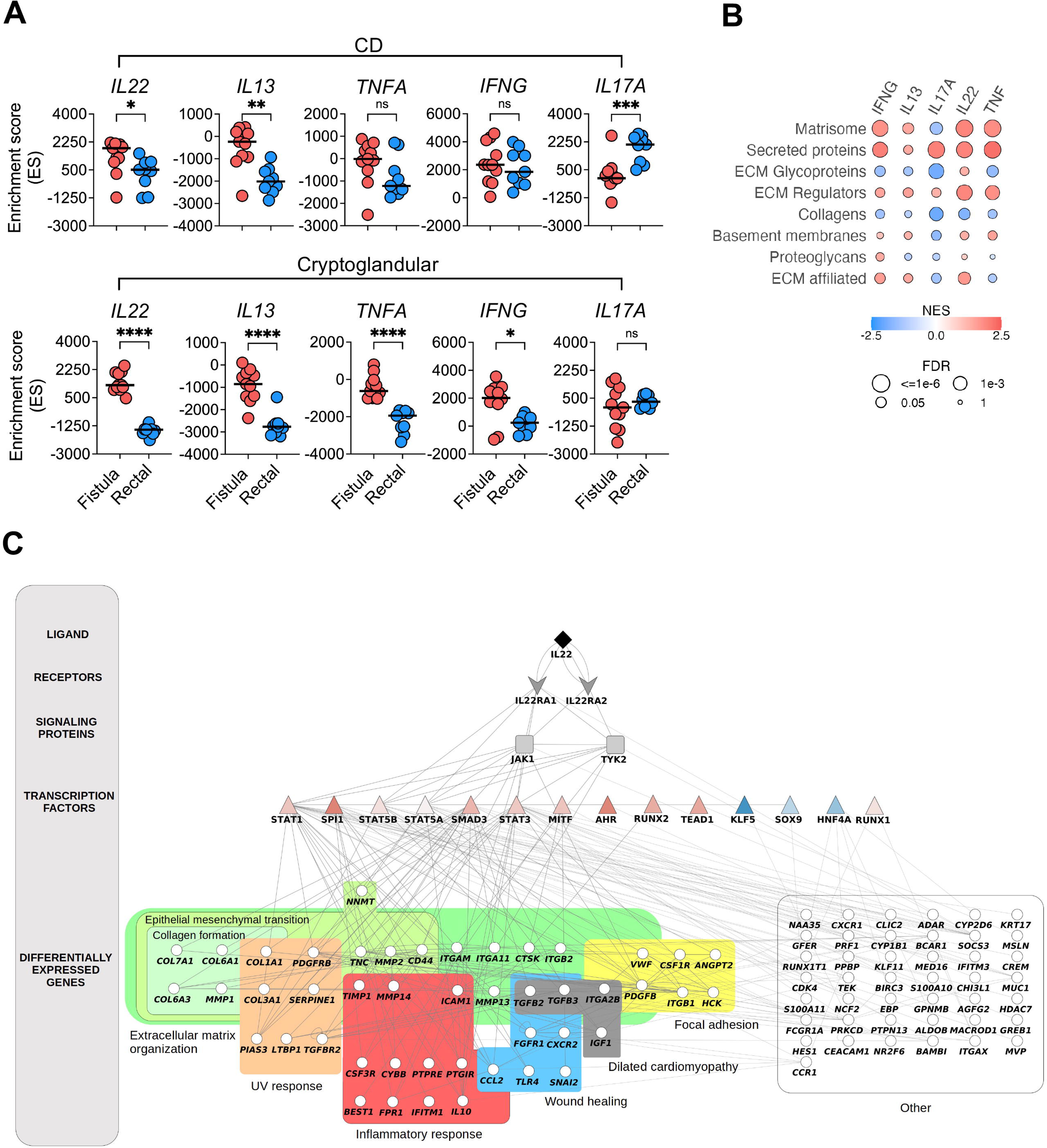
IL22 is a key regulator of the extracellular matrix and the matrisome. A) Distributions of single-sample GSEA scores for cytokine-responsive transcriptional signatures in fistula and rectal samples from patients with pCD or cryptoglandular disease. B) Bubble plot of GSEA results for matrisome-related gene sets in cytokine-treated colonic epithelial organoids. Circle fill colour and size represent the normalised enrichment score (NES) and false discovery rate (FDR) calculated by GSEA. C) Intracellular communication network showing interactions between IL22, components of the IL22 signalling pathway and impacted genes and pathways in perianal fistulas from pCD patients. Statistical significance of the differences in median values calculated by Mann-Whitney tests, \**P*<0.05, \*\**P*<0.01, \*\*\**P*<0.001, \*\*\*\**P*<0.0001,^ns^non-signifcant.

IL22 represents a potentially novel regulator of EMT and ECM remodelling in perianal fistulas. Given the prominent IL22 footprint at protein and gene-level in fistulas, we built a mechanistic model that might explain the role of IL22 in perianal fistulas and how it regulates gene expression changes between fistula and rectal tissue in pCD. Using the systems biology workflow ViralLink^30^, we constructed an intracellular communication network showing how IL22 regulated genes involved in EMT and the ECM in fistula tissue in pCD (**Figure 7C**). The network also included genes mapped to other disease relevant pathways including the inflammatory response, wound healing, focal adhesion, the UV response, and dilated cardiomyopathy. In addition, this visualization linked some highly activated transcription factors to the pathways identified by enrichment analysis. Collectively, the data presented in this paper indicate that canonical T-cell cytokines, particularly IL22 derived from CD4^+^ T-cells and iNKTs, are key modulators of EMT and tissue remodelling responses in pCD and cryptoglandular fistulas.

## Discussion

This study provides new insights into the immunopathology of pCD; implicating IL22 produced by conventional CD4^+^ T-cells and DN iNKT-cells as a potentially important driver of dysregulated matrisome function, a key biological process responsible for maintaining the balance between structural integrity of tissues and orderly remodelling following injury.

Conventional CD4^+^ T-cells and DN iNKT-cells were the dominant cellular producers of IL22 and IL13 in fistula tracts, and significantly accumulated in fistula tissue in comparison with peripheral blood in both pCD and cryptoglandular fistulas. Increased proportions of CD4^+^ T-cells and DN iNKT cells producing IL22 were most conspicuously observed in CD161 expressing subsets. The biological significance of CD161 expression is not well understood, but it has been linked to innate-like T-cells^35^, IL17 production^36^, and tissue residency^37–39^.

Tissue transcriptomics revealed activation of transcriptional programmes associated with EMT, ECM remodelling and innate and adaptive immune pathways, to be amongst the most upregulated in fistula tissue, which is broadly consistent with other studies^6,9,11,40^. This was further emphasized by the enrichment of genes encoding ECM proteins that form the “matrisome”^34^ and high expression of several proteases involved in tissue remodelling.

Overall, both flow cytometry and gene expression profiling of fistula tracts demonstrated considerable overlap in T-cell and molecular architecture of pCD and cryptoglandular fistulas. However, tissue transcriptomics was more discriminatory at uncovering molecular and biological pathway differences between the two clinical phenotypes. In a direct comparison of gene expression changes in biopsies from pCD and cryptoglandular fistula tracts, there was significant enrichment of pathways mapping to inflammation, cytokine signalling, T- and B-cell receptor signalling, JAK-STAT signalling, type I/II interferon responses, complement activation and innate immune activation in pCD. We also observed marked changes in the balance of MMPs and their natural inhibitors. In general, multiple protease families and MMPs were enriched in tissue of both pCD and cryptoglandular fistulas. However, the serine protease-inhibitor axis was preferentially dysregulated in pCD, with the expression of many cathepsin inhibitors conspicuously absent in pCD, but not cryptoglandular disease. Cathepsins are known to cleave ECM proteins and contribute to EMT and ECM remodelling during other related biological processes like angiogenesis^41^, cancer metastasis^42,43^ and collagen degradation in rheumatoid arthritis^44^.

Relatively little is known about the chief regulators of ECM and EMT in the fistula tract. Cytokines, such as TGFβ, TNFα and IL13 can induce expression of transcription factors linked to the control of cell invasion and EMT^6–8^, and now our work extends the potential molecular regulators of these critical processes to include IL22. IL22- responsive transcripts^25,26^ were more highly expressed in fistula tissue, including those associated with the ECM and its remodelling. Complementary network biology approaches also showed that IL22, through several key highly activated transcription factors, was responsible for regulating genes associated with EMT and wound healing in fistula tissue. It is conceptually attractive to envisage that IL22 derived from CD4^+^ T-cells and iNKT-cells act upon epithelial cells to drive EMT and ECM remodelling and promote fistula formation, propagation and persistence in pCD.

The implication of IL22 as a potentially important regulator of EMT and ECM remodelling in both pCD and cryptoglandular fistulas raises interesting questions. The function of IL22 is complex and is dependent upon cues from the surrounding tissue microenvironment. Under homeostatic conditions, IL22 contributes to epithelial repair and barrier maintenance^45,46^. However, in a chronic inflammatory setting it can trigger multiple proinflammatory pathways, including neutrophil recruitment through induction of CXC family chemokines^26^ and induction of endoplasmic reticulum stress in intestinal epithelial cells^47^.

Although IL22-responsive genes and IL22 producing cells were also enriched in cryptoglandular fistulas, and might potentially contribute to fistulization in this setting, it is also likely that the exaggerated pro-inflammatory tone of pCD fistulas alters the activity of IL22. The proinflammatory activities of IL22 are especially potent in synergy with other proinflammatory stimuli. For instance, induction of neutrophil-active chemokines and potentiation of intestinal epithelial apoptosis is augmented in the presence of other proinflammatory stimuli, including IL17A and TNFα^26,47^.

From a therapeutic perspective, identifying upstream drivers of IL22 and IL13 production by CD161^+^ CD4^+^ T-cells and CD161^+^ DN iNKT cells, or the key signalling pathways involved in their proximal activation and downstream effector function, are represent conceptually attractive targets in pCD. Notably, both IL13 and IL22 signal through JAK1, and selective inhibition of JAK1 with novel small molecule inhibitors may represent a novel way of simultaneously suppressing the inflammatory pathways and dysregulated tissue remodelling pathways mediated by these cytokines in fistulizing disease. Indeed, clinical studies evaluating JAK1 inhibition in pCD show promising results in preliminary or post hoc analyses^48,49^. Both conventional CD4^+^ T-cells and iNKT-cells are highly responsive to the microbiota, and in the case of iNKT-cells upon activation by microbial-derived lipids and glycosphingolipids, inducing production of proinflammatory cytokines^50–55^. This might at least partially explain the temporary beneficial effects seen following antibiotic treatment and highlights the promise of future strategies looking to modulate specific components of the microbiota, or their associated metabolites^56^.

In conclusion, fistulas remain a challenging complication of CD. Identification of IL22 producing cells, and augmented IL22-driven pathogenic pathways in Crohn’s tissue, highlights the importance of characterising the regulatory networks controlling the activation of IL22 producing lymphocytes, and the promise of therapeutically targeting this pathway in pCD.

## Author Contributions

Study concept and design conceived by N.P, A.H, P.T. Provided funding: N.P, A.H, P.T, T.K. Patient recruitment and consenting N.I. Data acquisition, analysis, or interpretation of data: L.E.C, D.C, L.C, N.P. Technical support: N.I, S.A, T.K. Drafting of the manuscript: L.E.C, N.P. Critical review of manuscript: all authors.

## Acknowledgements

We would also like to acknowledge the BRC FlowCore at King’s College London and the MRC LMS FlowCore facility at Imperial College London. We also acknowledge Novogene for performing the RNA-sequencing.

## References

1. Schwartz DA, Loftus EV, Tremaine WJ, et al. The natural history of fistulizing Crohn’s disease in Olmsted County, Minnesota. Gastroenterology 2002;122:875–880.

2. Adegbola SO, Dibley L, Sahnan K, et al. Burden of disease and adaptation to life in patients with Crohn’s perianal fistula: a qualitative exploration. Health Qual Life Outcomes 2020;18.

3. Stellingwerf ME, Praag EM van, Tozer PJ, et al. Systematic review and meta-analysis of endorectal advancement flap and ligation of the intersphincteric fistula tract for cryptoglandular and Crohn’s high perianal fistulas. BJS Open 2019;3:231–241.

4. Present DH, Rutgeerts P, Targan S, et al. Infliximab for the Treatment of Fistulas in Patients with Crohn’s Disease. New England Journal of Medicine 1999;340:1398–1405.

5. Sands BE, Anderson FH, Bernstein CN, et al. Infliximab Maintenance Therapy for Fistulizing Crohn’s Disease. New England Journal of Medicine 2004;350:876– 885.

6. Scharl M, Frei S, Pesch T, et al. Interleukin-13 and transforming growth factor β synergise in the pathogenesis of human intestinal fistulae. Gut 2013;62:63–72.

7. Frei SM, Pesch T, Lang S, et al. A Role for Tumor Necrosis Factor and Bacterial Antigens in the Pathogenesis of Crohn’s Disease–Associated Fistulae. Inflammatory Bowel Diseases 2013;19:2878–2887.

8. Bruckner RS, Spalinger MR, Barnhoorn MC, et al. Contribution of CD3+CD8- and CD3+CD8+ T Cells to TNF-α Overexpression in Crohn Disease-Associated Perianal Fistulas and Induction of Epithelial-Mesenchymal Transition in HT-29 Cells. Inflammatory Bowel Diseases 2021;27:538–549.

9. Bataille F, Klebl F, Rümmele P, et al. Morphological characterisation of Crohn’s disease fistulae. Gut 2004;53:1314–1321.

10. Bataille F, Rohrmeier C, Bates R, et al. Evidence for a role of epithelial mesenchymal transition during pathogenesis of fistulae in Crohn’s disease. Inflammatory Bowel Diseases 2008;14:1514–1527.

11. Scharl M, Weber A, Fürst A, et al. Potential role for SNAIL family transcription factors in the etiology of Crohn’s disease-associated fistulae. Inflammatory Bowel Diseases 2011;17:1907–1916.

12. Kirkegaard T, Hansen A, Bruun E, et al. Expression and localisation of matrix metalloproteinases and their natural inhibitors in fistulae of patients with Crohn’s disease. Gut 2004;53:701–709.

13. Goffin L, Fagagnini S, Vicari A, et al. Anti-MMP-9 Antibody: A Promising Therapeutic Strategy for Treatment of Inflammatory Bowel Disease Complications with Fibrosis. Inflammatory Bowel Diseases 2016;22:2041–2057.

14. Maggi L, Capone M, Giudici F, et al. CD4+CD161+ T lymphocytes infiltrate Crohn’s disease-associated perianal fistulas and are reduced by anti-TNF-α local therapy. Int Arch Allergy Immunol 2013;161:81–86.

15. Bolger AM, Lohse M, Usadel B. Trimmomatic: a flexible trimmer for Illumina sequence data. Bioinformatics 2014;30:2114–2120.

16. Kim D, Langmead B, Salzberg SL. HISAT: a fast spliced aligner with low memory requirements. Nat Methods 2015;12:357–360.

17. Howe KL, Achuthan P, Allen J, et al. Ensembl 2021. Nucleic Acids Research 2021;49:D884–D891.

18. Anders S, Pyl PT, Huber W. HTSeq--a Python framework to work with high-throughput sequencing data. Bioinformatics 2015;31:166–169.

19. Core R. Team. R: A Language and Environment for Statistical Computing, 2015. 2021.

20. Love MI, Huber W, Anders S. Moderated estimation of fold change and dispersion for RNA-seq data with DESeq2. Genome Biology 2014;15:550.

21. Hochberg Y, Benjamini Y. More powerful procedures for multiple significance testing. Stat Med 1990;9:811–818.

22. Liberzon A, Birger C, Thorvaldsdóttir H, et al. The Molecular Signatures Database (MSigDB) hallmark gene set collection. Cell Syst 2015;1:417–425.

23. Liberzon A, Subramanian A, Pinchback R, et al. Molecular signatures database (MSigDB) 3.0. Bioinformatics 2011;27:1739–1740.

24. Subramanian A, Tamayo P, Mootha VK, et al. Gene set enrichment analysis: A knowledge-based approach for interpreting genome-wide expression profiles. PNAS 2005;102:15545–15550.

25. Pavlidis P, Tsakmaki A, Treveil A, et al. Cytokine responsive networks in human colonic epithelial organoids unveil a molecular classification of inflammatory bowel disease. Cell Reports 2022;40.

26. Pavlidis P, Tsakmaki A, Pantazi E, et al. Interleukin-22 regulates neutrophil recruitment in ulcerative colitis and is associated with resistance to ustekinumab therapy. Nat Commun 2022;13:5820.

27. Hänzelmann S, Castelo R, Guinney J. GSVA: gene set variation analysis for microarray and RNA-Seq data. BMC Bioinformatics 2013;14:7.

28. Alvarez MJ, Shen Y, Giorgi FM, et al. Functional characterization of somatic mutations in cancer using network-based inference of protein activity. Nat Genet 2016;48:838–847.

29. Garcia-Alonso L, Holland CH, Ibrahim MM, et al. Benchmark and integration of resources for the estimation of human transcription factor activities. Genome Res 2019;29:1363–1375.

30. Treveil A, Bohar B, Sudhakar P, et al. ViralLink: An integrated workflow to investigate the effect of SARS-CoV-2 on intracellular signalling and regulatory pathways. PLOS Computational Biology 2021;17:e1008685.

31. Anon. Gene Set - il22 soluble receptor signaling pathway. Available at: https://maayanlab.cloud/Harmonizome/gene_set/il22+soluble+receptor+signaling+pathway/Biocarta+Pathways [Accessed March 13, 2023].

32. Shannon P, Markiel A, Ozier O, et al. Cytoscape: a software environment for integrated models of biomolecular interaction networks. Genome Res 2003;13:2498–2504.

33. Rovedatti L, Kudo T, Biancheri P, et al. Differential regulation of interleukin 17 and interferon gamma production in inflammatory bowel disease. Gut 2009;58:1629–1636.

34. Hynes RO, Naba A. Overview of the Matrisome—An Inventory of Extracellular Matrix Constituents and Functions. Cold Spring Harb Perspect Biol 2012;4:a004903.

35. Fergusson JR, Smith KE, Fleming VM, et al. CD161 Defines a Transcriptional and Functional Phenotype across Distinct Human T Cell Lineages. Cell Reports 2014;9:1075–1088.

36. Maggi L, Santarlasci V, Capone M, et al. CD161 is a marker of all human IL-17- producing T-cell subsets and is induced by RORC. European Journal of Immunology 2010;40:2174–2181.

37. Billerbeck E, Kang Y-H, Walker L, et al. Analysis of CD161 expression on human CD8+ T cells defines a distinct functional subset with tissue-homing properties. PNAS 2010;107:3006–3011.

38. Fergusson JR, Hühn MH, Swadling L, et al. CD161intCD8+ T cells: a novel population of highly functional, memory CD8+ T cells enriched within the gut. Mucosal Immunology 2016;9:401–413.

39. Takahashi T, Dejbakhsh-Jones S, Strober S. Expression of CD161 (NKR-P1A) defines subsets of human CD4 and CD8 T cells with different functional activities. J Immunol 2006;176:211–216.

40. Rizzo G, Rubbino F, Elangovan S, et al. Dysfunctional Extracellular Matrix Remodeling Supports Perianal Fistulizing Crohn’s Disease by a Mechanoregulated Activation of the Epithelial-to-Mesenchymal Transition. Cell Mol Gastroenterol Hepatol 2023;15:741–764.

41. Nakao S, Zandi S, Sun D, et al. Cathepsin B-mediated CD18 shedding regulates leukocyte recruitment from angiogenic vessels. FASEB J 2018;32:143–154.

42. Wang B, Sun J, Kitamoto S, et al. Cathepsin S controls angiogenesis and tumor growth via matrix-derived angiogenic factors. J Biol Chem 2006;281:6020–6029.

43. Podgorski I, Linebaugh BE, Koblinski JE, et al. Bone Marrow-Derived Cathepsin K Cleaves SPARC in Bone Metastasis. The American Journal of Pathology 2009;175:1255–1269.

44. Mort JS, Beaudry F, Théroux K, et al. Early cathepsin K degradation of type II collagen in vitro and in vivo in articular cartilage. Osteoarthritis Cartilage 2016;24:1461–1469.

45. Lindemans CA, Calafiore M, Mertelsmann AM, et al. Interleukin-22 promotes intestinal-stem-cell-mediated epithelial regeneration. Nature 2015;528:560–564.

46. Aparicio-Domingo P, Romera-Hernandez M, Karrich JJ, et al. Type 3 innate lymphoid cells maintain intestinal epithelial stem cells after tissue damage. J Exp Med 2015;212:1783–1791.

47. Powell N, Pantazi E, Pavlidis P, et al. Interleukin-22 orchestrates a pathological endoplasmic reticulum stress response transcriptional programme in colonic epithelial cells. Gut 2020;69:578–590.

48. Reinisch W, Colombel JF, D’Haens GR, et al. OP18 Efficacy and safety of filgotinib for the treatment of perianal fistulizing Crohn’s Disease: Results from the phase 2 DIVERGENCE 2 study. Journal of Crohn’s and Colitis 2022;16:i019– i021.

49. Colombel JF, Irving P, Rieder F, et al. P491 Efficacy and safety of upadacitinib for the treatment of fistulas and fissures in patients with Crohn’s disease. Journal of Crohn’s and Colitis 2023;17:i620–i623.

50. Lantz O, Bendelac A. An invariant T cell receptor alpha chain is used by a unique subset of major histocompatibility complex class I-specific CD4+ and CD4-8- T cells in mice and humans. J Exp Med 1994;180:1097–1106.

51. Borg NA, Wun KS, Kjer-Nielsen L, et al. CD1d–lipid-antigen recognition by the semi-invariant NKT T-cell receptor. Nature 2007;448:44–49.

52. Kawano T, Cui J, Koezuka Y, et al. CD1d-Restricted and TCR-Mediated Activation of Vα14 NKT Cells by Glycosylceramides. Science 1997;278:1626– 1629.

53. Exley M, Garcia J, Balk SP, et al. Requirements for CD1d Recognition by Human Invariant Vα24+ CD4−CD8− T Cells. Journal of Experimental Medicine 1997;186:109–120.

54. Burrello C, Pellegrino G, Giuffrè MR, et al. Mucosa-associated microbiota drives pathogenic functions in IBD-derived intestinal iNKT cells. Life Sci Alliance 2019;2.

55. Adegbola SO, Sarafian M, Sahnan K, et al. Differences in amino acid and lipid metabolism distinguish Crohn’s from idiopathic/cryptoglandular perianal fistulas by tissue metabonomic profiling and may offer clues to underlying pathogenesis. European Journal of Gastroenterology & Hepatology 2021;33:1469–1479.

56. Cao S, Colonna M, Deepak P. Pathogenesis of Perianal Fistulising Crohn’s Disease: Current Knowledge, Gaps in Understanding, and Future Research Directions. Journal of Crohn’s and Colitis 2023;17:1010–1022.

