## Supplementary material for "Invariant natural killer T-cell and CD4^+^ T-cell derived IL22 is a regulator of epithelial-to-mesenchymal transition and extracellular matrix remodelling in perianal fistulas"

**Tables**

Supplementary Table 1. Characteristics of patients with CD (n=14) or cryptoglandular (n=17) perianal fistulizing disease from who peripheral blood and fistula curettage samples were collected and used during immune phenotyping experiments. Age, HBI, PDAI and FDA were presented as mean ± SD. Other variables were presented as a percentage within each group.

| | **CD (n=14)** | **Cryptoglandular (n=17)** |
|---|---|---|
| **Age (years)** | 36.5 (15.5) | 48.6 (13.3) |
| **Gender** | | |
| Male | 6/14 (43%) | 11/17 (65%) |
| Female | 8/14 (57%) | 6/17 (35%) |
| **CD duration (years)** | | |
| 0-5 | 2/14 (14%) | n/a |
| 5-7 | 1/14 (7%) | n/a |
| >7 | 10/14 (71%) | n/a |
| Unknown | 1/14 (7%) | n/a |
| **Fistula duration (years)** | | |
| 0-2 | 4/14 (29%) | 9/17 (53%) |
| 2-4 | 1/14 (7%) | 2/17 (12%) |
| 4-12 | 7/14 (50%) | 5/17 (29%) |
| 12-24 | 1/14 (7%) | 0/17 (0%) |
| >24 | 1/14 (7%) | 0/17 (0%) |
| Unknown | 0/14 (0%) | 1/17 (6%) |
| **Concomitant medications** | | |
| Anti-TNF | 3/14 (21%) | n/a |
| Anti-TNF & immunomodulators | 7/14 (50%) | n/a |
| Anti-TNF & 5-ASAs | 1/14 (7%) | n/a |
| Other biologics (e.g., Vedolizumab) | 1/14 (7%) | n/a |
| None | 2/14 (14%) | n/a |
| **Harvey Bradshaw Index (HBI)** | 1.2 (1.2) | n/a |
| **Perianal disease activity index (PDAI)** | 8.1 (1.8) | 9.6 (2.9) |
| **Fistula drainage assessment (FDA)** | 1.3 (0.5) | 1.7 (2.2) |
| **Park's classification** | | |
| Intersphincteric | 1 (7%) | 2/17 (12%) |
| Transphincteric | 9 (64%) | 12/17 (71%) |
| Extrasphincteric | 1 (7%) | 0/17 (0%) |
| Suprasphincteric | 1 (7%) | 0/17 (0%) |
| Intersphincteric and transphincteric | 1 (7%) | 1/17 (6%) |
| Transphincteric and extrasphincteric | 1 (7%) | 0/17 (0%) |

| | | |
|---|---|---|
| Intersphincteric and suprasphincteric | 0 (0%) | 1/17 (6%) |
| Unknown | 0 (0%) | 1/17 (6%) |
| **Number of internal openings** | | |
| 0 | 0/14 (0%) | 0/17 (0%) |
| 1 | 11/14 (79%) | 12/17 (71%) |
| 2 | 3/14 (21%) | 5/17 (29%) |
| **Number of external openings** | | |
| 0 | 0/14 (0%) | 1/17 (6%) |
| 1 | 10/14 (71%) | 12/17 (71%) |
| 2 | 4/14 (29%) | 3/17 (17%) |
| >2 | 0/14 (0%) | 1/17 (6%) |
| **Morphological features** | | |
| Horseshoe | 2/14 (14%) | 5/17 (29%) |
| Cavities | 4/14 (29%) | 3/17 (17%) |
| **Number of branches** | | |
| 0 | 9/14 (64%) | 7/17 (41%) |
| 1 | 3/14 (21%) | 7/17 (41%) |
| 2 | 1/14 (7%) | 2/17 (12%) |
| Unknown | 1/14 (7%) | 1/17 (6%) |
| **Active luminal disease** | 1/14 (7%) | n/a |
| **Proctitis** | 1/14 (7%) | n/a |

Supplementary Table 2. Characteristics of patients with CD (n=13) or cryptoglandular (n=13) perianal fistulizing disease from who fistula and rectal biopsies were collected and used during RNA-sequencing experiments. Age, HBI, PDAI and FDA were presented as mean ± SD. Other variables were presented as a percentage within each group.

| | **CD (n=13)** | **Cryptoglandular (n=13)** |
|---|---|---|
| **Age (years)** | 33.8 (15.1) | 47.6 (12.9) |
| **Gender** | | |
| Male | 5/13 (38%) | 7/13 (54%) |
| Female | 8/13 (62%) | 6/13 (46%) |
| **CD duration (years)** | | |
| 0-5 | 3/13 (23%) | n/a |
| 5-7 | 0/13 (0%) | n/a |
| >7 | 9/13 (69%) | n/a |
| Unknown | 1/13 (8%) | n/a |
| **Fistula duration (years)** | | |
| 0-2 | 3/13 (23%) | 7/13 (54%) |
| 2-4 | 1/13 (8%) | 2/13 (15%) |
| 4-12 | 6/13 (46%) | 3/13 (23%) |
| 12-24 | 1/13 (8%) | 0/13 (0%) |
| >24 | 1/13 (8%) | 0/13 (0%) |
| Unknown | 1/13 (8%) | 1/13 (8%) |
| **Concomitant medications** | | |
| Anti-TNF | 4/13 (31%) | n/a |
| Anti-TNF & immunomodulators | 8/13 (62%) | n/a |
| Anti-TNF & 5-ASAs | 1/13 (8%) | n/a |
| Other biologics (e.g., Vedolizumab) | 0/13 (0%) | n/a |
| None | 0/13 (0%) | n/a |
| **Harvey Bradshaw Index (HBI)** | 1.6 (2.1) | n/a |
| **Perianal disease activity index (PDAI)** | 8.9 (2.4) | 9.4 (3.1) |
| **Fistula drainage assessment (FDA)** | 1.3 (0.5) | 1.2 (0.4) |
| **Park's classification** | | |
| Intersphincteric | 0/13 (0%) | 1/13 (8%) |
| Transphincteric | 9/13 (69%) | 10/13 (77%) |
| Extrasphincteric | 1/13 (8%) | 0/13 (0%) |
| Suprasphincteric | 0/13 (0%) | 0/13 (0%) |
| Intersphincteric and transphincteric | 2/13 (15%) | 1/13 (8%) |
| Transphincteric and extrasphincteric | 1/13 (8%) | 0/13 (0%) |
| Intersphincteric and suprasphincteric | 0/13 (0%) | 0/13 (0%) |
| Unknown | 0/13 (0%) | 1/13 (8%) |
| **Number of internal openings** | | |
| 0 | 0/13 (0%) | 0/13 (0%) |

| | | |
|---|---|---|
| 1 | 9/13 (69%) | 10/13 (77%) |
| 2 | 4/14 (31%) | 3/13 (23%) |
| **Number of external openings** | | |
| 0 | 0/13 (0%) | 0/13 (0%) |
| 1 | 9/13 (69%) | 11/13 (65%) |
| 2 | 4/13 (31%) | 2/13 (15%) |
| >2 | 0/13 (0%) | 0/13 (0%) |
| **Morphological features** | | |
| Horseshoe | 2/13 (15%) | 4/13 (31%) |
| Cavities | 2/13 (15%) | 2/13 (15%) |
| **Number of branches** | | |
| 0 | 8/13 (62%) | 6/13 (46%) |
| 1 | 4/13 (31%) | 4/13 (31%) |
| 2 | 1/13 (8%) | 2/13 (15%) |
| Unknown | 0/13 (0%) | 1/13 (8%) |
| **Active luminal disease** | 1/13 (8%) | n/a |
| **Proctitis** | 1/13 (8%) | n/a |

Supplementary Table 3. Assignment of biological pathways to clusters identified by hierarchical clustering if impacted pathways identified by gene set enrichment analysis in fistula versus rectal tissue in CD and cryptoglandular fistulizing disease.

| **Pathway** | **Cluster assignment** |
|---|---|
| HALLMARK_ANGIOGENESIS | 1 |
| HALLMARK_APICAL_JUNCTION | 1 |
| HALLMARK_COAGULATION | 1 |
| HALLMARK_E2F_TARGETS | 1 |
| HALLMARK_EPITHELIAL_MESENCHYMAL_TRANSITION | 1 |
| HALLMARK_ESTROGEN_RESPONSE_EARLY | 1 |
| HALLMARK_ESTROGEN_RESPONSE_LATE | 1 |
| HALLMARK_FATTY_ACID_METABOLISM | 1 |
| HALLMARK_G2M_CHECKPOINT | 1 |
| HALLMARK_INFLAMMATORY_RESPONSE | 1 |
| HALLMARK_MYOGENESIS | 1 |
| HALLMARK_PANCREAS_BETA_CELLS | 1 |
| HALLMARK_PEROXISOME | 1 |
| HALLMARK_UV_RESPONSE_DN | 1 |
| KEGG_BUTANOATE_METABOLISM | 1 |
| KEGG_CYTOKINE_CYTOKINE_RECEPTOR_INTERACTION | 1 |
| KEGG_DILATED_CARDIOMYOPATHY | 1 |
| KEGG_GLYCOLYSIS_GLUCONEOGENESIS | 1 |
| KEGG_GLYCOSPHINGOLIPID_BIOSYNTHESIS_LACTO_AND_NEOLACTO_SERIES | 1 |
| KEGG_HEMATOPOIETIC_CELL_LINEAGE | 1 |
| KEGG_HYPERTROPHIC_CARDIOMYOPATHY_HCM | 1 |
| KEGG_MATURITY_ONSET_DIABETES_OF_THE_YOUNG | 1 |
| KEGG_PEROXISOME | 1 |
| KEGG_RIBOSOME | 1 |
| PID_AMB2_NEUTROPHILS_PATHWAY | 1 |
| PID_AR_TF_PATHWAY | 1 |
| PID_FRA_PATHWAY | 1 |
| PID_HNF3B_PATHWAY | 1 |
| PID_INTEGRIN_CS_PATHWAY | 1 |
| PID_LYMPH_ANGIOGENESIS_PATHWAY | 1 |
| PID_S1P_S1P1_PATHWAY | 1 |

| | |
|---|---|
| PID_TOLL_ENDOGENOUS_PATHWAY | 1 |
| PID_UPA_UPAR_PATHWAY | 1 |
| REACTOME_ACTIVATED_PKN1_STIMULATES_TRANSCRIPTION_OF_AR_ANDROGEN_RECEPTOR_REGULATED_GENES_KLK2_AND_KLK3 | 1 |
| REACTOME_ANTI_INFLAMMATORY_RESPONSE_FAVOURING_LEISHMANIA_PARASITE_INFECTION | 1 |
| REACTOME_BLOOD_GROUP_SYSTEMS_BIOSYNTHESIS | 1 |
| REACTOME_BUDDING_AND_MATURATION_OF_HIV_VIRION | 1 |
| REACTOME_C_TYPE_LECTIN_RECEPTORS_CLRS | 1 |
| REACTOME_CRISTAE_FORMATION | 1 |
| REACTOME_CRMPS_IN_SEMA3A_SIGNALING | 1 |
| REACTOME_CROSSLINKING_OF_COLLAGEN_FIBRILS | 1 |
| REACTOME_DIGESTION | 1 |
| REACTOME_DIGESTION_AND_ABSORPTION | 1 |
| REACTOME_DISEASES_OF_GLYCOSYLATION | 1 |
| REACTOME_ELASTIC_FIBRE_FORMATION | 1 |
| REACTOME_EUKARYOTIC_TRANSLATION_ELONGATION | 1 |
| REACTOME_FATTY_ACID_METABOLISM | 1 |
| REACTOME_FORMATION_OF_THE_CORNIFIED_ENVELOPE | 1 |
| REACTOME_G_ALPHA_Z_SIGNALLING_EVENTS | 1 |
| REACTOME_GABA_B_RECEPTOR_ACTIVATION | 1 |
| REACTOME_GLUCONEOGENESIS | 1 |
| REACTOME_GLYCEROPHOSPHOLIPID_BIOSYNTHESIS | 1 |
| REACTOME_GPCR_LIGAND_BINDING | 1 |
| REACTOME_INCRETIN_SYNTHESIS_SECRETION_AND_INACTIVATION | 1 |
| REACTOME_INTERLEUKIN_4_AND_INTERLEUKIN_13_SIGNALING | 1 |
| REACTOME_INWARDLY_RECTIFYING_K_CHANNELS | 1 |
| REACTOME_ION_CHANNEL_TRANSPORT | 1 |
| REACTOME_KERATINIZATION | 1 |
| REACTOME_LEISHMANIA_INFECTION | 1 |
| REACTOME_MITOCHONDRIAL_FATTY_ACID_BETA_OXIDATION | 1 |
| REACTOME_MITOCHONDRIAL_TRANSLATION | 1 |
| REACTOME_MOLECULES_ASSOCIATED_WITH_ELASTIC_FIBRES | 1 |
| REACTOME_NOTCH4_INTRACELLULAR_DOMAIN_REGULATES_TRANSCRIPTION | 1 |
| REACTOME_NUCLEAR_RECEPTOR_TRANSCRIPTION_PATHWAY | 1 |

| | |
|---|---|
| REACTOME_O_GLYCOSYLATION_OF_TSR_DOMAIN_CONTAINING_PROTEINS | 1 |
| REACTOME_PEROXISOMAL_LIPID_METABOLISM | 1 |
| REACTOME_PEROXISOMAL_PROTEIN_IMPORT | 1 |
| REACTOME_PHOSPHOLIPID_METABOLISM | 1 |
| REACTOME_PLATELET_ACTIVATION_SIGNALING_AND_AGGREGATION | 1 |
| REACTOME_PLATELET_ADHESION_TO_EXPOSED_COLLAGEN | 1 |
| REACTOME_PLATELET_AGGREGATION_PLUG_FORMATION | 1 |
| REACTOME_PROTEIN_LOCALIZATION | 1 |
| REACTOME_PROTEIN_PROTEIN_INTERACTIONS_AT_SYNAPSES | 1 |
| REACTOME_REGULATION_OF_BETA_CELL_DEVELOPMENT | 1 |
| REACTOME_REGULATION_OF_GENE_EXPRESSION_IN_BETA_CELLS | 1 |
| REACTOME_REGULATION_OF_INSULIN_LIKE_GROWTH_FACTOR_IGF_TRANSPORT_AND_UPTAKE_BY_INSULIN_LIKE_GROWTH_FACTOR_BINDING_PROTEINS_IGFBPS | 1 |
| REACTOME_RESPONSE_TO_ELEVATED_PLATELET_CYTOSOLIC_CA2 | 1 |
| REACTOME_SCAVENGING_BY_CLASS_A_RECEPTORS | 1 |
| REACTOME_SIGNALING_BY_VEGF | 1 |
| REACTOME_SLC_MEDIATED_TRANSMEMBRANE_TRANSPORT | 1 |
| REACTOME_SLC_TRANSPORTER_DISORDERS | 1 |
| REACTOME_SMOOTH_MUSCLE_CONTRACTION | 1 |
| REACTOME_STIMULI_SENSING_CHANNELS | 1 |
| REACTOME_SYNTHESIS_OF_PA | 1 |
| REACTOME_SYNTHESIS_OF_SUBSTRATES_IN_N_GLYCAN_BIOSYTHESIS | 1 |
| REACTOME_SYNTHESIS_SECRETION_AND_INACTIVATION_OF_GLUCAGON_LIKE_PEPTIDE_1_GLP_1 | 1 |
| REACTOME_TIGHT_JUNCTION_INTERACTIONS | 1 |
| REACTOME_TRANSLATION | 1 |
| REACTOME_TRANSPORT_OF_BILE_SALTS_AND_ORGANIC_ACIDS_METAL_IONS_AND_AMINE_COMPOUNDS | 1 |
| WP_ANGIOGENESIS | 1 |
| WP_BURN_WOUND_HEALING | 1 |
| WP_CANONICAL_AND_NONCANONICAL_NOTCH_SIGNALING | 1 |
| WP_CCL18_SIGNALING_PATHWAY | 1 |
| WP_CILIOPATHIES | 1 |

| | |
|---|---|
| WP_COMPLEMENT_SYSTEM | 1 |
| WP_CONSTITUTIVE_ANDROSTANE_RECEPTOR_PATHWAY | 1 |
| WP_FAMILIAL_HYPERLIPIDEMIA_TYPE_1 | 1 |
| WP_FAMILIAL_HYPERLIPIDEMIA_TYPE_4 | 1 |
| WP_FOXA2_PATHWAY | 1 |
| WP_GDNFRET_SIGNALING_AXIS | 1 |
| WP_GPCRS_CLASS_A_RHODOPSINLIKE | 1 |
| WP_GPCRS_OTHER | 1 |
| WP_HAIR_FOLLICLE_DEVELOPMENT_CYTODIFFERENTIATION_PART_3_OF_3 | 1 |
| WP_HIPPOMERLIN_SIGNALING_DYSREGULATION | 1 |
| WP_HYPOTHESIZED_PATHWAYS_IN_PATHOGENESIS_OF_CARDIOVASCULAR_DISEASE | 1 |
| WP_INFLAMMATORY_RESPONSE_PATHWAY | 1 |
| WP_MIR5093P_ALTERATION_OF_YAP1ECM_AXIS | 1 |
| WP_MIRNA_TARGETS_IN_ECM_AND_MEMBRANE_RECEPTORS | 1 |
| WP_MITOCHONDRIAL_COMPLEX_IV_ASSEMBLY | 1 |
| WP_NEOVASCULARISATION_PROCESSES | 1 |
| WP_NETRINUNC5B_SIGNALING_PATHWAY | 1 |
| WP_NETWORK_MAP_OF_SARSCOV2_SIGNALING_PATHWAY | 1 |
| WP_NRF2_PATHWAY | 1 |
| WP_NRP1TRIGGERED_SIGNALING_PATHWAYS_IN_PANCREATIC_CANCER | 1 |
| WP_NUCLEAR_RECEPTORS_METAPATHWAY | 1 |
| WP_OREXIN_RECEPTOR_PATHWAY | 1 |
| WP_PHOTODYNAMIC_THERAPYINDUCED_NFKB_SURVIVAL_SIGNALING | 1 |
| WP_PLATELETMEDIATED_INTERACTIONS_WITH_VASCULAR_AND_CIRCULATING_CELLS | 1 |
| WP_PREGNANE_X_RECEPTOR_PATHWAY | 1 |
| WP_PROXIMAL_TUBULE_TRANSPORT | 1 |
| WP_SMALL_LIGAND_GPCRS | 1 |
| WP_SPINAL_CORD_INJURY | 1 |
| WP_TYPE_I_COLLAGEN_SYNTHESIS_IN_THE_CONTEXT_OF_OSTEOGENESIS_IMPERFECTA | 1 |
| WP_ZINC_HOMEOSTASIS | 1 |
| PID_SYNDECAN_1_PATHWAY | 2 |
| REACTOME_ASSEMBLY_OF_COLLAGEN_FIBRILS_AND_OTHER_MULTIMERIC_STRUCTURES | 2 |
| REACTOME_COLLAGEN_BIOSYNTHESIS_AND_MODIFYING_ENZYMES | 2 |

| | |
|---|---|
| REACTOME_COLLAGEN_CHAIN_TRIMERIZATION | 2 |
| REACTOME_COLLAGEN_DEGRADATION | 2 |
| REACTOME_COLLAGEN_FORMATION | 2 |
| REACTOME_DEGRADATION_OF_THE_EXTRACELLULAR_MATRIX | 2 |
| REACTOME_EXTRACELLULAR_MATRIX_ORGANIZATION | 2 |
| REACTOME_NCAM_SIGNALING_FOR_NEURITE_OUT_GROWTH | 2 |
| REACTOME_NCAM1_INTERACTIONS | 2 |
| REACTOME_SIGNALING_BY_PDGF | 2 |
| HALLMARK_OXIDATIVE_PHOSPHORYLATION | 3 |
| KEGG_ALZHEIMERS_DISEASE | 3 |
| KEGG_HUNTINGTONS_DISEASE | 3 |
| KEGG_OXIDATIVE_PHOSPHORYLATION | 3 |
| KEGG_PARKINSONS_DISEASE | 3 |
| REACTOME_COMPLEX_I_BIOGENESIS | 3 |
| REACTOME_RESPIRATORY_ELECTRON_TRANSPORT | 3 |
| REACTOME_RESPIRATORY_ELECTRON_TRANSPORT_ATP_SYNTHESIS_BY_CHEMIOSMOTIC_COUPLING_AND_HEAT_PRODUCTION_BY_UNCOUPLING_PROTEINS | 3 |
| REACTOME_THE_CITRIC_ACID_TCA_CYCLE_AND_RESPIRATORY_ELECTRON_TRANSPORT | 3 |
| WP_ELECTRON_TRANSPORT_CHAIN_OXPHOS_SYSTEM_IN_MITOCHONDRIA | 3 |
| WP_MITOCHONDRIAL_COMPLEX_I_ASSEMBLY_MODEL_OXPHOS_SYSTEM | 3 |
| WP_NONALCOHOLIC_FATTY_LIVER_DISEASE | 3 |
| WP_OXIDATIVE_PHOSPHORYLATION | 3 |
| KEGG_ECM_RECEPTOR_INTERACTION | 4 |
| KEGG_FOCAL_ADHESION | 4 |
| PID_AVB3_INTEGRIN_PATHWAY | 4 |
| PID_INTEGRIN1_PATHWAY | 4 |
| PID_INTEGRIN3_PATHWAY | 4 |
| REACTOME_ECM_PROTEOGLYCANS | 4 |
| REACTOME_INTEGRIN_CELL_SURFACE_INTERACTIONS | 4 |
| REACTOME_LAMININ_INTERACTIONS | 4 |
| REACTOME_MET_ACTIVATES_PTK2_SIGNALING | 4 |
| REACTOME_MET_PROMOTES_CELL_MOTILITY | 4 |
| REACTOME_NON_INTEGRIN_MEMBRANE_ECM_INTERACTIONS | 4 |
| REACTOME_SIGNALING_BY_MET | 4 |
| REACTOME_SYNDECAN_INTERACTIONS | 4 |

| | |
|---|---|
| WP_FOCAL_ADHESION | 4 |
| WP_FOCAL_ADHESION_PI3KAKTMTORSIGNALING_PATHWAY | 4 |
| WP_PI3KAKT_SIGNALING_PATHWAY | 4 |
| REACTOME_ANTIGEN_ACTIVATES_B_CELL_RECEPTOR_BCR_LEADING_TO_GENERATION_OF_SECOND_MESSENGERS | 5 |
| REACTOME_BINDING_AND_UPTAKE_OF_LIGANDS_BY_SCAVENGER_RECEPTORS | 5 |
| REACTOME_CD22_MEDIATED_BCR_REGULATION | 5 |
| REACTOME_CELL_SURFACE_INTERACTIONS_AT_THE_VASCULAR_WALL | 5 |
| REACTOME_COMPLEMENT_CASCADE | 5 |
| REACTOME_CREATION_OF_C4_AND_C2_ACTIVATORS | 5 |
| REACTOME_FCERI_MEDIATED_CA_2_MOBILIZATION | 5 |
| REACTOME_FCGAMMA_RECEPTOR_FCGR_DEPENDENT_PHAGOCYTOSIS | 5 |
| REACTOME_FCGR_ACTIVATION | 5 |
| REACTOME_FCGR3A_MEDIATED_IL10_SYNTHESIS | 5 |
| REACTOME_IMMUNOREGULATORY_INTERACTIONS_BETWEEN_A_LYMPHOID_AND_A_NON_LYMPHOID_CELL | 5 |
| REACTOME_INITIAL_TRIGGERING_OF_COMPLEMENT | 5 |
| REACTOME_PARASITE_INFECTION | 5 |
| REACTOME_ROLE_OF_LAT2_NTAL_LAB_ON_CALCIUM_MOBILIZATION | 5 |
| REACTOME_ROLE_OF_PHOSPHOLIPIDS_IN_PHAGOCYTOSIS | 5 |
| REACTOME_SCAVENGING_OF_HEME_FROM_PLASMA | 5 |
| KEGG_GLYCOSAMINOGLYCAN_BIOSYNTHESIS_CHONDROITIN_SULFATE | 6 |
| REACTOME_CHONDROITIN_SULFATE_BIOSYNTHESIS | 6 |
| REACTOME_CHONDROITIN_SULFATE_DERMATAN_SULFATE_METABOLISM | 6 |
| REACTOME_CS_DS_DEGRADATION | 6 |
| REACTOME_DISEASES_ASSOCIATED_WITH_GLYCOSAMINOGLYCAN_METABOLISM | 6 |
| REACTOME_GLYCOSAMINOGLYCAN_METABOLISM | 6 |
| KEGG_ASCORBATE_AND_ALDARATE_METABOLISM | 7 |
| KEGG_DRUG_METABOLISM_OTHER_ENZYMES | 7 |
| KEGG_METABOLISM_OF_XENOBIOTICS_BY_CYTOCHROME_P450 | 7 |
| KEGG_PENTOSE_AND_GLUCURONATE_INTERCONVERSIONS | 7 |
| KEGG_PORPHYRIN_AND_CHLOROPHYLL_METABOLISM | 7 |
| KEGG_RETINOL_METABOLISM | 7 |

| | |
|---|---|
| KEGG_STARCH_AND_SUCROSE_METABOLISM | 7 |
| KEGG_STEROID_HORMONE_BIOSYNTHESIS | 7 |
| REACTOME_BIOLOGICAL_OXIDATIONS | 7 |
| REACTOME_GLUCURONIDATION | 7 |
| REACTOME_PHASE_II_CONJUGATION_OF_COMPOUNDS | 7 |
| WP_GLUCURONIDATION | 7 |
| REACTOME_DECTIN_2_FAMILY | 8 |
| REACTOME_DEFECTIVE_C1GALT1C1_CAUSES_TNPS | 8 |
| REACTOME_DEFECTIVE_GALNT3_CAUSES_HFTC | 8 |
| REACTOME_O_LINKED_GLYCOSYLATION_OF_MUCINS | 8 |
| REACTOME_TERMINATION_OF_O_GLYCAN_BIOSYNTHESIS | 8 |
| WP_BREAST_CANCER_PATHWAY | 9 |
| WP_LNCRNA_IN_CANONICAL_WNT_SIGNALING_AND_COLORECTAL_CANCER | 9 |
| WP_NCRNAS_INVOLVED_IN_WNT_SIGNALING_IN_HEPATOCELLULAR_CARCINOMA | 9 |
| WP_WNT_SIGNALING | 9 |
| KEGG_CITRATE_CYCLE_TCA_CYCLE | 10 |
| REACTOME_CITRIC_ACID_CYCLE_TCA_CYCLE | 10 |
| REACTOME_PYRUVATE_METABOLISM_AND_CITRIC_ACID_TCA_CYCLE | 10 |
| WP_TCA_CYCLE_AKA_KREBS_OR_CITRIC_ACID_CYCLE | 10 |

Supplementary Table 4. Assignment of biological pathways to clusters identified by hierarchical clustering of impacted pathways identified by gene set enrichment analysis in CD versus cryptoglandular fistula tissue.

| Pathway | Cluster assignment |
|---|---|
| REACTOME_FORMATION_OF_THE_CORNIFIED_ENVELOPE | 1 |
| REACTOME_KERATINIZATION | 1 |
| WP_HAIR_FOLLICLE_DEVELOPMENT_CYTODIFFERENTIATION_PART_3_OF_3 | 1 |
| REACTOME_TRANSLATION | 1 |
| HALLMARK_ESTROGEN_RESPONSE_EARLY | 1 |
| WP_NUCLEAR_RECEPTORS_METAPATHWAY | 1 |
| HALLMARK_ESTROGEN_RESPONSE_LATE | 1 |
| KEGG_CELL_ADHESION_MOLECULES_CAMS | 1 |
| REACTOME_SPHINGOLIPID_DE_NOVO_BIOSYNTHESIS | 1 |
| HALLMARK_P53_PATHWAY | 1 |
| WP_EPITHELIAL_TO_MESENCHYMAL_TRANSITION_IN_COLORECTAL_CANCER | 1 |
| REACTOME_SPHINGOLIPID_METABOLISM | 1 |
| WP_OXIDATION_BY_CYTOCHROME_P450 | 1 |
| REACTOME_GAP_JUNCTION_TRAFFICKING_AND_REGULATION | 1 |
| REACTOME_APOPTOTIC_EXECUTION_PHASE | 1 |
| REACTOME_RND3_GTPASE_CYCLE | 1 |
| REACTOME_APOPTOTIC_CLEAVAGE_OF_CELL_ADHESION_PROTEINS | 1 |
| REACTOME_SLC_MEDIATED_TRANSMEMBRANE_TRANSPORT | 1 |
| WP_GLUCOCORTICOID_RECEPTOR_PATHWAY | 1 |
| REACTOME_SENSORY_PERCEPTION | 1 |
| REACTOME_SYNTHESIS_OF_SUBSTRATES_IN_N_GLYCAN_BIOSYTHESIS | 1 |
| WP_NRF2_PATHWAY | 1 |
| REACTOME_TRANSPORT_OF_BILE_SALTS_AND_ORGANIC_ACIDS_METAL_IONS_AND_AMINE_COMPOUNDS | 1 |
| WP_HEAD_AND_NECK_SQUAMOUS_CELL_CARCINOMA | 1 |
| REACTOME_GAP_JUNCTION_ASSEMBLY | 1 |
| REACTOME_APOPTOTIC_CLEAVAGE_OF_CELLULAR_PROTEINS | 1 |
| WP_WNT_SIGNALING_PATHWAY_AND_PLURIPOTENCY | 1 |
| REACTOME_CELL_CELL_JUNCTION_ORGANIZATION | 1 |

| | |
|---|---|
| PID_BETA_CATENIN_NUC_PATHWAY | 1 |
| REACTOME_RND2_GTPASE_CYCLE | 1 |
| WP_SPHINGOLIPID_PATHWAY | 1 |
| REACTOME_MITOCHONDRIAL_TRANSLATION | 1 |
| REACTOME_VISUAL_PHOTOTRANSDUCTION | 1 |
| WP_SPHINGOLIPID_METABOLISM_OVERVIEW | 1 |
| KEGG_STEROID_HORMONE_BIOSYNTHESIS | 1 |
| KEGG_TIGHT_JUNCTION | 1 |
| REACTOME_BIOSYNTHESIS_OF_THE_N_GLYCAN_PRECURSOR_DOLICHOL_LIPID_LINKED_OLIGOSACCHARIDE_LLO_AND_TRANSFER_TO_A_NASCENT_PROTEIN | 1 |
| PID_TAP63_PATHWAY | 1 |
| REACTOME_REGULATION_OF_TP53_ACTIVITY_THROUGH_ASSOCIATION_WITH_CO_FACTORS | 1 |
| REACTOME_DEFECTIVE_INTRINSIC_PATHWAY_FOR_APOPTOSIS | 1 |
| REACTOME_THE_CANONICAL_RETINOID_CYCLE_IN_RODS_TWILIGHT_VISION | 1 |
| WP_HAIR_FOLLICLE_DEVELOPMENT_ORGANOGENESIS_PART_2_OF_3 | 1 |
| REACTOME_TP53_REGULATES_TRANSCRIPTION_OF_GENES_INVOLVED_IN_CYTOCHROME_C_RELEASE | 1 |
| PID_FOXO_PATHWAY | 1 |
| REACTOME_CELL_JUNCTION_ORGANIZATION | 1 |
| PID_DELTA_NP63_PATHWAY | 1 |
| WP_VITAMIN_A_AND_CAROTENOID_METABOLISM | 1 |
| WP_METAPATHWAY_BIOTRANSFORMATION_PHASE_I_AND_II | 1 |
| REACTOME_AMINO_ACID_TRANSPORT_ACROSS_THE_PLASMA_MEMBRANE | 1 |
| REACTOME_TIGHT_JUNCTION_INTERACTIONS | 1 |
| PID_MTOR_4PATHWAY | 1 |
| HALLMARK_ALLOGRAFT_REJECTION | 2 |
| HALLMARK_INTERFERON_GAMMA_RESPONSE | 2 |
| HALLMARK_INTERFERON_ALPHA_RESPONSE | 2 |
| KEGG_CYTOKINE_CYTOKINE_RECEPTOR_INTERACTION | 2 |
| PID_IL12_2PATHWAY | 2 |
| KEGG_HEMATOPOIETIC_CELL_LINEAGE | 2 |
| REACTOME_COSTIMULATION_BY_THE_CD28_FAMILY | 2 |
| KEGG_ANTIGEN_PROCESSING_AND_PRESENTATION | 2 |
| WP_MICROGLIA_PATHOGEN_PHAGOCYTOSIS_PATHWAY | 2 |
| KEGG_PRIMARY_IMMUNODEFICIENCY | 2 |
| REACTOME_INTERLEUKIN_2_FAMILY_SIGNALING | 2 |
| WP_INFLAMMATORY_RESPONSE_PATHWAY | 2 |
| WP_NETWORK_MAP_OF_SARSCOV2_SIGNALING_PATHWAY | 2 |

| | |
|---|---|
| REACTOME_GENERATION_OF_SECOND_MESSENGER_MOLECULES | 2 |
| KEGG_VIRAL_MYOCARDITIS | 2 |
| HALLMARK_KRAS_SIGNALING_DN | 2 |
| WP_INTERACTIONS_BETWEEN_IMMUNE_CELLS_AND_MICRORNAS_IN_TUMOR_MICROENVIRONMENT | 2 |
| REACTOME_INTERFERON_ALPHA_BETA_SIGNALING | 2 |
| REACTOME_PD_1_SIGNALING | 2 |
| WP_TYROBP_CAUSAL_NETWORK_IN_MICROGLIA | 2 |
| WP_PATHOGENESIS_OF_SARSCOV2_MEDIATED_BY_NSP9NSP10_COMPLEX | 2 |
| PID_IL12_STAT4_PATHWAY | 2 |
| WP_TYPE_II_INTERFERON_SIGNALING_IFNG | 2 |
| WP_SELECTIVE_EXPRESSION_OF_CHEMOKINE_RECEPTORS_DURING_TCELL_POLARIZATION | 2 |
| REACTOME_TNF_RECEPTOR_SUPERFAMILY_TNFSF_MEMBERS_MEDIATING_NON_CANONICAL_NF_KB_PATHWAY | 2 |
| KEGG_NATURAL_KILLER_CELL_MEDIATED_CYTOTOXICITY | 2 |
| WP_CANCER_IMMUNOTHERAPY_BY_PD1_BLOCKADE | 2 |
| HALLMARK_IL6_JAK_STAT3_SIGNALING | 2 |
| PID_CD8_TCR_DOWNSTREAM_PATHWAY | 2 |
| WP_GENES_ASSOCIATED_WITH_THE_DEVELOPMENT_OF_RHEUMATOID_ARTHRITIS | 2 |
| REACTOME_INTERFERON_SIGNALING | 2 |
| WP_FOXP3_IN_COVID19 | 2 |
| WP_PLATELETMEDIATED_INTERACTIONS_WITH_VASCULAR_AND_CIRCULATING_CELLS | 2 |
| REACTOME_INTERLEUKIN_2_SIGNALING | 2 |
| PID_CXCR4_PATHWAY | 2 |
| HALLMARK_HYPOXIA | 2 |
| HALLMARK_TNFA_SIGNALING_VIA_NFKB | 2 |
| REACTOME_SIGNAL_REGULATORY_PROTEIN_FAMILY_INTERACTIONS | 2 |
| WP_EBOLA_VIRUS_INFECTION_IN_HOST | 2 |
| PID_IL4_2PATHWAY | 2 |
| WP_COMPLEMENT_SYSTEM | 2 |
| WP_ACUTE_VIRAL_MYOCARDITIS | 2 |
| HALLMARK_COMPLEMENT | 2 |
| REACTOME_TNFS_BIND_THEIR_PHYSIOLOGICAL_RECEPTORS | 2 |
| KEGG_JAK_STAT_SIGNALING_PATHWAY | 2 |
| WP_COMPLEMENT_AND_COAGULATION_CASCADES | 2 |
| REACTOME_INTERLEUKIN_RECEPTOR_SHC_SIGNALING | 2 |
| PID_INTEGRIN2_PATHWAY | 2 |
| WP_IMMUNE_RESPONSE_TO_TUBERCULOSIS | 2 |
| WP_PREIMPLANTATION_EMBRYO | 2 |

| | |
|---|---|
| REACTOME_TRANSCRIPTIONAL_REGULATION_BY_THE_AP_2_TFAP2_FAMILY_OF_TRANSCRIPTION_FACTORS | 2 |
| REACTOME_INTERFERON_GAMMA_SIGNALING | 2 |
| REACTOME_INACTIVATION_OF_CSF3_G_CSF_SIGNALING | 2 |
| HALLMARK_IL2_STAT5_SIGNALING | 2 |
| KEGG_COMPLEMENT_AND_COAGULATION_CASCADES | 2 |
| REACTOME_ANTIGEN_PRESENTATION_FOLDING_ASSEMBLY_AND_PEPTIDE_LOADING_OF_CLASS_I_MHC | 2 |
| REACTOME_THE_ROLE_OF_NEF_IN_HIV_1_REPLICATION_AND_DISEASE_PATHOGENESIS | 2 |
| REACTOME_INTERLEUKIN_3_INTERLEUKIN_5_AND_GM_CSF_SIGNALING | 2 |
| PID_IL2_STAT5_PATHWAY | 2 |
| HALLMARK_INFLAMMATORY_RESPONSE | 2 |
| REACTOME_IMMUNOREGULATORY_INTERACTIONS_BETWEEN_A_LYMPHOID_AND_A_NON_LYMPHOID_CELL | 3 |
| REACTOME_COMPLEMENT_CASCADE | 3 |
| REACTOME_SCAVENGING_OF_HEME_FROM_PLASMA | 3 |
| REACTOME_CD22_MEDIATED_BCR_REGULATION | 3 |
| REACTOME_INITIAL_TRIGGERING_OF_COMPLEMENT | 3 |
| REACTOME_ANTIGEN_ACTIVATES_B_CELL_RECEPTOR_BCR_LEADING_TO_GENERATION_OF_SECOND_MESSENGERS | 3 |
| REACTOME_CREATION_OF_C4_AND_C2_ACTIVATORS | 3 |
| REACTOME_CELL_SURFACE_INTERACTIONS_AT_THE_VASCULAR_WALL | 3 |
| REACTOME_FCGR_ACTIVATION | 3 |
| REACTOME_BINDING_AND_UPTAKE_OF_LIGANDS_BY_SCAVENGER_RECEPTORS | 3 |
| REACTOME_ROLE_OF_LAT2_NTAL_LAB_ON_CALCIUM_MOBILIZATION | 3 |
| REACTOME_FCERI_MEDIATED_CA_2_MOBILIZATION | 3 |
| REACTOME_PARASITE_INFECTION | 3 |
| REACTOME_ROLE_OF_PHOSPHOLIPIDS_IN_PHAGOCYTOSIS | 3 |
| REACTOME_LEISHMANIA_INFECTION | 3 |
| REACTOME_FCGAMMA_RECEPTOR_FCGR_DEPENDENT_PHAGOCYTOSIS | 3 |
| REACTOME_FCERI_MEDIATED_MAPK_ACTIVATION | 3 |
| REACTOME_ANTI_INFLAMMATORY_RESPONSE_FAVOURING_LEISHMANIA_PARASITE_INFECTION | 3 |
| REACTOME_FCGR3A_MEDIATED_IL10_SYNTHESIS | 3 |
| REACTOME_SRP_DEPENDENT_COTRANSLATIONAL_PROTEIN_TARGETING_TO_MEMBRANE | 4 |
| KEGG_RIBOSOME | 4 |

| | |
|---|---|
| REACTOME_EUKARYOTIC_TRANSLATION_ELONGATION | 4 |
| WP_CYTOPLASMIC_RIBOSOMAL_PROTEINS | 4 |
| REACTOME_RESPONSE_OF_EIF2AK4_GCN2_TO_AMINO_ACID_DEFICIENCY | 4 |
| REACTOME_SELENOAMINO_ACID_METABOLISM | 4 |
| REACTOME_NONSENSE_MEDIATED_DECAY_NMD | 4 |
| REACTOME_EUKARYOTIC_TRANSLATION_INITIATION | 4 |
| REACTOME_REGULATION_OF_EXPRESSION_OF_SLITS_AND_ROBOS | 4 |
| REACTOME_CELLULAR_RESPONSE_TO_STARVATION | 4 |
| REACTOME_INFLUENZA_INFECTION | 4 |
| REACTOME_SIGNALING_BY_ROBO_RECEPTORS | 4 |
| REACTOME_RRNA_PROCESSING | 4 |
| REACTOME_ACTIVATION_OF_THE_MRNA_UPON_BINDING_OF_THE_CAP_BINDING_COMPLEX_AND_EIFS_AND_SUBSEQUENT_BINDING_TO_43S | 4 |
| KEGG_INTESTINAL_IMMUNE_NETWORK_FOR_IGA_PRODUCTION | 5 |
| WP_ALLOGRAFT_REJECTION | 5 |
| KEGG_AUTOIMMUNE_THYROID_DISEASE | 5 |
| KEGG_ALLOGRAFT_REJECTION | 5 |
| KEGG_SYSTEMIC_LUPUS_ERYTHEMATOSUS | 5 |
| KEGG_TYPE_I_DIABETES_MELLITUS | 5 |
| KEGG_LEISHMANIA_INFECTION | 5 |
| KEGG_ASTHMA | 5 |
| KEGG_GRAFT_VERSUS_HOST_DISEASE | 5 |
| PID_TCR_PATHWAY | 6 |
| PID_CD8_TCR_PATHWAY | 6 |
| WP_MODULATORS_OF_TCR_SIGNALING_AND_T_CELL_ACTIVATION | 6 |
| WP_TCELL_ACTIVATION_SARSCOV2 | 6 |
| WP_TCELL_RECEPTOR_TCR_SIGNALING_PATHWAY | 6 |
| WP_TCELL_ANTIGEN_RECEPTOR_TCR_PATHWAY_DURING_STAPHYLOCOCCUS_AUREUS_INFECTION | 6 |
| KEGG_T_CELL_RECEPTOR_SIGNALING_PATHWAY | 6 |
| WP_B_CELL_RECEPTOR_SIGNALING_PATHWAY | 6 |
| REACTOME_SIGNALING_BY_THE_B_CELL_RECEPTOR_BCR | 7 |
| REACTOME_TCR_SIGNALING | 7 |
| REACTOME_FCERI_MEDIATED_NF_KB_ACTIVATION | 7 |
| REACTOME_TNFR2_NON_CANONICAL_NF_KB_PATHWAY | 7 |
| REACTOME_FC_EPSILON_RECEPTOR_FCERI_SIGNALING | 7 |
| REACTOME_ANTIGEN_PROCESSING_CROSS_PRESENTATION | 7 |
| WP_PROTEASOME_DEGRADATION | 7 |

| | |
|---|---|
| REACTOME_CHEMOKINE_RECEPTORS_BIND_CHEMOKINES | 8 |
| REACTOME_CLASS_A_1_RHODOPSIN_LIKE_RECEPTORS | 8 |
| KEGG_CHEMOKINE_SIGNALING_PATHWAY | 8 |
| REACTOME_G_ALPHA_I_SIGNALLING_EVENTS | 8 |
| REACTOME_PEPTIDE_LIGAND_BINDING_RECEPTORS | 8 |
| WP_CHEMOKINE_SIGNALING_PATHWAY | 8 |
| WP_PEPTIDE_GPCRS | 8 |
| REACTOME_GPCR_LIGAND_BINDING | 8 |
| WP_GPCRS_CLASS_A_RHODOPSINLIKE | 8 |
| REACTOME_RESPIRATORY_ELECTRON_TRANSPORT_ATP_SYNTHESIS_BY_CHEMIOSMOTIC_COUPLING_AND_HEAT_PRODUCTION_BY_UNCOUPLING_PROTEINS | 9 |
| WP_ELECTRON_TRANSPORT_CHAIN_OXPHOS_SYSTEM_IN_MITOCHONDRIA | 9 |
| REACTOME_RESPIRATORY_ELECTRON_TRANSPORT | 9 |
| REACTOME_COMPLEX_I_BIOGENESIS | 9 |
| WP_MITOCHONDRIAL_COMPLEX_I_ASSEMBLY_MODEL_OXPHOS_SYSTEM | 9 |
| KEGG_PARKINSONS_DISEASE | 9 |
| REACTOME_METABOLISM_OF_STEROIDS | 10 |
| REACTOME_REGULATION_OF_CHOLESTEROL_BIOSYNTHESIS_BY_SREBP_SREBF | 10 |
| REACTOME_ACTIVATION_OF_GENE_EXPRESSION_BY_SREBF_SREBP | 10 |
| REACTOME_CHOLESTEROL_BIOSYNTHESIS | 10 |
| WP_CHOLESTEROL_METABOLISM_WITH_BLOCH_AND_KANDUTSCHRUSSELL_PATHWAYS | 10 |
| WP_CHOLESTEROL_BIOSYNTHESIS_PATHWAY | 10 |

Supplementary Table 5. Assignment of biological pathways to clusters identified by hierarchical clustering of impacted pathways identified by gene set enrichment analysis in CD and cryptoglandular rectal tissue.

| **Pathway** | **Cluster assignment** |
|---|---|
| REACTOME_SRP_DEPENDENT_COTRANSLATIONAL_PROTEIN_TARGETING_TO_MEMBRANE | 1 |
| WP_CYTOPLASMIC_RIBOSOMAL_PROTEINS | 1 |
| REACTOME_EUKARYOTIC_TRANSLATION_ELONGATION | 1 |
| REACTOME_RESPONSE_OF_EIF2AK4_GCN2_TO_AMINO_ACID_DEFICIENCY | 1 |
| KEGG_RIBOSOME | 1 |
| REACTOME_SELENOAMINO_ACID_METABOLISM | 1 |
| REACTOME_EUKARYOTIC_TRANSLATION_INITIATION | 1 |
| REACTOME_CELLULAR_RESPONSE_TO_STARVATION | 1 |
| REACTOME_NONSENSE_MEDIATED_DECAY_NMD | 1 |
| REACTOME_REGULATION_OF_EXPRESSION_OF_SLITS_AND_ROBOS | 1 |
| REACTOME_INFLUENZA_INFECTION | 1 |
| REACTOME_RRNA_PROCESSING | 1 |
| REACTOME_SIGNALING_BY_ROBO_RECEPTORS | 1 |
| REACTOME_METABOLISM_OF_AMINO_ACIDS_AND_DERIVATIVES | 1 |
| REACTOME_ACTIVATION_OF_THE_MRNA_UPON_BINDING_OF_THE_CAP_BINDING_COMPLEX_AND_EIFS_AND_SUBSEQUENT_BINDING_TO_43S | 1 |
| REACTOME_IMMUNOREGULATORY_INTERACTIONS_BETWEEN_A_LYMPHOID_AND_A_NON_LYMPHOID_CELL | 2 |
| REACTOME_COMPLEMENT_CASCADE | 2 |
| REACTOME_INITIAL_TRIGGERING_OF_COMPLEMENT | 2 |
| REACTOME_CREATION_OF_C4_AND_C2_ACTIVATORS | 2 |
| REACTOME_ANTIGEN_ACTIVATES_B_CELL_RECEPTOR_BCR_LEADING_TO_GENERATION_OF_SECOND_MESSENGERS | 2 |
| REACTOME_CELL_SURFACE_INTERACTIONS_AT_THE_VASCULAR_WALL | 2 |
| REACTOME_SCAVENGING_OF_HEME_FROM_PLASMA | 2 |
| REACTOME_CD22_MEDIATED_BCR_REGULATION | 2 |
| REACTOME_FCGR_ACTIVATION | 2 |

| | |
|---|---|
| REACTOME_SIGNALING_BY_THE_B_CELL_RECEPTOR_BCR | 2 |
| REACTOME_FCERI_MEDIATED_CA_2_MOBILIZATION | 2 |
| REACTOME_FCERI_MEDIATED_NF_KB_ACTIVATION | 2 |
| REACTOME_ROLE_OF_PHOSPHOLIPIDS_IN_PHAGOCYTOSIS | 2 |
| REACTOME_BINDING_AND_UPTAKE_OF_LIGANDS_BY_SCAVENGER_RECEPTORS | 2 |
| REACTOME_ROLE_OF_LAT2_NTAL_LAB_ON_CALCIUM_MOBILIZATION | 2 |
| REACTOME_FCGR3A_MEDIATED_IL10_SYNTHESIS | 2 |
| REACTOME_FCERI_MEDIATED_MAPK_ACTIVATION | 2 |
| REACTOME_FC_EPSILON_RECEPTOR_FCERI_SIGNALING | 2 |
| REACTOME_PARASITE_INFECTION | 2 |
| REACTOME_ANTI_INFLAMMATORY_RESPONSE_FAVOURING_LEISHMANIA_PARASITE_INFECTION | 2 |
| REACTOME_FCGAMMA_RECEPTOR_FCGR_DEPENDENT_PHAGOCYTOSIS | 2 |
| REACTOME_LEISHMANIA_INFECTION | 2 |
| REACTOME_TRANSLATION | 3 |
| HALLMARK_ALLOGRAFT_REJECTION | 3 |
| HALLMARK_E2F_TARGETS | 3 |
| KEGG_SYSTEMIC_LUPUS_ERYTHEMATOSUS | 3 |
| HALLMARK_PROTEIN_SECRETION | 3 |
| KEGG_HEMATOPOIETIC_CELL_LINEAGE | 3 |
| HALLMARK_MYC_TARGETS_V1 | 3 |
| WP_TYROBP_CAUSAL_NETWORK_IN_MICROGLIA | 3 |
| HALLMARK_G2M_CHECKPOINT | 3 |
| REACTOME_SLC_MEDIATED_TRANSMEMBRANE_TRANSPORT | 3 |
| WP_COMPLEMENT_SYSTEM | 3 |
| REACTOME_TNFS_BIND_THEIR_PHYSIOLOGICAL_RECEPTORS | 3 |
| HALLMARK_HEME_METABOLISM | 3 |
| REACTOME_MITOCHONDRIAL_TRANSLATION | 3 |
| WP_NUCLEAR_RECEPTORS_IN_LIPID_METABOLISM_AND_TOXICITY | 3 |
| REACTOME_SPHINGOLIPID_METABOLISM | 3 |
| HALLMARK_IL6_JAK_STAT3_SIGNALING | 3 |
| REACTOME_ION_TRANSPORT_BY_P_TYPE_ATPASES | 3 |
| REACTOME_INTERLEUKIN_4_AND_INTERLEUKIN_13_SIGNALING | 3 |

| | |
|---|---|
| REACTOME_TRANS_GOLGI_NETWORK_VESICLE_BUDDING | 3 |
| HALLMARK_ANDROGEN_RESPONSE | 3 |
| REACTOME_INTRA_GOLGI_AND_RETROGRADE_GOLGI_TO_ER_TRAFFIC | 3 |
| WP_STEROL_REGULATORY_ELEMENTBINDING_PROTEINS_SREBP_SIGNALING | 3 |
| REACTOME_ANTIMICROBIAL_PEPTIDES | 3 |
| REACTOME_RETROGRADE_TRANSPORT_AT_THE_TRANS_GOLGI_NETWORK | 3 |
| HALLMARK_MYC_TARGETS_V2 | 3 |
| PID_IL4_2PATHWAY | 3 |
| KEGG_COMPLEMENT_AND_COAGULATION_CASCADES | 3 |
| WP_RETINOBLASTOMA_GENE_IN_CANCER | 3 |
| REACTOME_DISASSEMBLY_OF_THE_DESTRUCTION_COMPLEX_AND_RECRUITMENT_OF_AXIN_TO_THE_MEMBRANE | 3 |
| REACTOME_GOLGI_ASSOCIATED_VESICLE_BIOGENESIS | 3 |
| WP_NUCLEAR_RECEPTORS | 3 |
| REACTOME_SIGNALING_BY_WNT_IN_CANCER | 3 |
| KEGG_DNA_REPLICATION | 3 |
| PID_INTEGRIN2_PATHWAY | 3 |
| REACTOME_NUCLEAR_RECEPTOR_TRANSCRIPTION_PATHWAY | 3 |
| HALLMARK_TGF_BETA_SIGNALING | 3 |
| REACTOME_SPHINGOLIPID_DE_NOVO_BIOSYNTHESIS | 3 |
| REACTOME_TRANSPORT_OF_BILE_SALTS_AND_ORGANIC_ACIDS_METAL_IONS_AND_AMINE_COMPOUNDS | 3 |
| REACTOME_REGULATION_OF_LIPID_METABOLISM_BY_PPARALPHA | 3 |
| KEGG_SPHINGOLIPID_METABOLISM | 3 |
| HALLMARK_INFLAMMATORY_RESPONSE | 3 |
| WP_16P112_DISTAL_DELETION_SYNDROME | 3 |
| REACTOME_DNA_STRAND_ELONGATION | 3 |
| REACTOME_O_LINKED_GLYCOSYLATION_OF_MUCINS | 3 |
| REACTOME_APC_C_CDC20_MEDIATED_DEGRADATION_OF_CYCLIN_B | 3 |
| WP_DNA_REPLICATION | 3 |
| KEGG_VIRAL_MYOCARDITIS | 3 |
| KEGG_O_GLYCAN_BIOSYNTHESIS | 3 |
| WP_NUCLEAR_RECEPTORS_METAPATHWAY | 3 |

| | |
|---|---|
| WP_COMPLEMENT_ACTIVATION | 3 |
| PID_FOXO_PATHWAY | 3 |
| WP_COMPLEMENT_AND_COAGULATION_CASCADES | 3 |
| REACTOME_CHROMOSOME_MAINTENANCE | 3 |
| WP_PREGNANE_X_RECEPTOR_PATHWAY | 3 |
| WP_HEMATOPOIETIC_STEM_CELL_DIFFERENTIATION | 3 |
| REACTOME_HDMS_DEMETHYLATE_HISTONES | 3 |
| HALLMARK_DNA_REPAIR | 3 |
| KEGG_ENDOCYTOSIS | 3 |
| KEGG_STARCH_AND_SUCROSE_METABOLISM | 3 |
| WP_CONSTITUTIVE_ANDROSTANE_RECEPTOR_PATHWAY | 3 |
| WP_TGFBETA_SIGNALING_PATHWAY | 3 |
| REACTOME_APC_CDC20_MEDIATED_DEGRADATION_OF_NEK2A | 3 |
| WP_1Q211_COPY_NUMBER_VARIATION_SYNDROME | 3 |
| REACTOME_RAB_GEFS_EXCHANGE_GTP_FOR_GDP_ON_RABS | 3 |
| REACTOME_FOXO_MEDIATED_TRANSCRIPTION | 3 |
| REACTOME_SIGNALING_BY_CTNNB1_PHOSPHO_SITE_MUTANTS | 3 |
| REACTOME_O_LINKED_GLYCOSYLATION | 3 |
| REACTOME_PHOSPHOLIPID_METABOLISM | 3 |
| KEGG_CIRCADIAN_RHYTHM_MAMMAL | 3 |
| WP_COMPLEMENT_SYSTEM_IN_NEURONAL_DEVELOPMENT_AND_PLASTICITY | 3 |
| WP_ANDROGEN_RECEPTOR_SIGNALING_PATHWAY | 3 |
| HALLMARK_IL2_STAT5_SIGNALING | 3 |
| REACTOME_PHASE_I_FUNCTIONALIZATION_OF_COMPOUNDS | 3 |
| WP_OXIDATION_BY_CYTOCHROME_P450 | 3 |
| WP_GENES_ASSOCIATED_WITH_THE_DEVELOPMENT_OF_RHEUMATOID_ARTHRITIS | 3 |
| HALLMARK_UV_RESPONSE_DN | 3 |
| REACTOME_COPI_INDEPENDENT_GOLGI_TO_ER_RETROGRADE_TRAFFIC | 3 |
| REACTOME_TRANSPORT_TO_THE_GOLGI_AND_SUBSEQUENT_MODIFICATION | 3 |
| PID_BETA_CATENIN_DEG_PATHWAY | 3 |
| REACTOME_ACTIVATION_OF_THE_PRE_REPLICATIVE_COMPLEX | 3 |
| WP_INTERACTIONS_BETWEEN_IMMUNE_CELLS_AND_MICRORNAS_IN_TUMOR_MICROENVIRONMENT | 3 |

| | |
|---|---|
| REACTOME_BETA_CATENIN_PHOSPHORYLATION_CASCADE | 3 |
| REACTOME_UNWINDING_OF_DNA | 3 |
| PID_HIF2PATHWAY | 3 |
| REACTOME_SENSORY_PERCEPTION | 3 |
| REACTOME_ION_CHANNEL_TRANSPORT | 3 |
| WP_PROXIMAL_TUBULE_TRANSPORT | 3 |
| WP_INFLAMMATORY_RESPONSE_PATHWAY | 3 |
| WP_SPHINGOLIPID_PATHWAY | 3 |
| HALLMARK_COMPLEMENT | 3 |
| PID_FOXM1_PATHWAY | 3 |
| KEGG_ABC_TRANSPORTERS | 3 |
| REACTOME_GLYCEROPHOSPHOLIPID_BIOSYNTHESIS | 3 |
| WP_DEVELOPMENT_OF_PULMONARY_DENDRITIC_CELLS_AND_MACROPHAGE_SUBSETS | 3 |
| REACTOME_MISCELLANEOUS_TRANSPORT_AND_BINDING_EVENTS | 3 |
| REACTOME_MITOCHONDRIAL_PROTEIN_IMPORT | 3 |
| HALLMARK_MITOTIC_SPINDLE | 3 |
| KEGG_ASCORBATE_AND_ALDARATE_METABOLISM | 3 |
| WP_ACUTE_VIRAL_MYOCARDITIS | 3 |
| PID_NFAT_TFPATHWAY | 3 |
| REACTOME_HSP90_CHAPERONE_CYCLE_FOR_STEROID_HORMONE_RECEPTORS_SHR_IN_THE_PRESENCE_OF_LIGAND | 3 |
| REACTOME_CARNITINE_METABOLISM | 3 |
| REACTOME_SLC_TRANSPORTER_DISORDERS | 3 |
| KEGG_CELL_CYCLE | 3 |
| REACTOME_KERATINIZATION | 3 |
| REACTOME_GLUCURONIDATION | 3 |
| REACTOME_TNF_RECEPTOR_SUPERFAMILY_TNFSF_MEMBERS_MEDIATING_NON_CANONICAL_NF_KB_PATHWAY | 3 |
| WP_COPPER_HOMEOSTASIS | 3 |
| REACTOME_PHOSPHORYLATION_OF_THE_APC_C | 3 |
| PID_AMB2_NEUTROPHILS_PATHWAY | 3 |
| REACTOME_ACTIVATION_OF_ATR_IN_RESPONSE_TO_REPLICATION_STRESS | 3 |
| WP_THERMOGENESIS | 3 |
| PID_WNT_CANONICAL_PATHWAY | 3 |
| WP_MELATONIN_METABOLISM_AND_EFFECTS | 3 |
| REACTOME_COHESIN_LOADING_ONTO_CHROMATIN | 3 |

| | |
|---|---|
| PID_TXA2PATHWAY | 3 |
| WP_BIOMARKERS_FOR_PYRIMIDINE_METABOLISM_DISORDERS | 3 |
| WP_PRION_DISEASE_PATHWAY | 3 |
| REACTOME_REGULATION_OF_CHOLESTEROL_BIOSYNTHESIS_BY_SREBP_SREBF | 3 |
| REACTOME_UPTAKE_AND_ACTIONS_OF_BACTERIAL_TOXINS | 3 |
| WP_CELL_CYCLE | 3 |
| REACTOME_ER_TO_GOLGI_ANTEROGRADE_TRANSPORT | 3 |
| REACTOME_FORMATION_OF_THE_CORNIFIED_ENVELOPE | 3 |
| WP_MATRIX_METALLOPROTEINASES | 3 |
| REACTOME_INFLAMMASOMES | 3 |
| WP_PRIMARY_FOCAL_SEGMENTAL_GLOMERULOSCLEROSIS_FSGS | 3 |
| KEGG_RETINOL_METABOLISM | 3 |
| HALLMARK_ESTROGEN_RESPONSE_EARLY | 3 |
| REACTOME_TRANSLOCATION_OF_SLC2A4_GLUT4_TO_THE_PLASMA_MEMBRANE | 3 |
| WP_SREBF_AND_MIR33_IN_CHOLESTEROL_AND_LIPID_HOMEOSTASIS | 3 |
| REACTOME_APOPTOTIC_CLEAVAGE_OF_CELLULAR_PROTEINS | 3 |
| WP_ARYL_HYDROCARBON_RECEPTOR_PATHWAY_WP2586 | 3 |
| REACTOME_SUPPRESSION_OF_PHAGOSOMAL_MATURATION | 3 |
| PID_AR_TF_PATHWAY | 3 |
| REACTOME_REGULATION_OF_FOXO_TRANSCRIPTIONAL_ACTIVITY_BY_ACETYLATION | 3 |
| PID_TGFBR_PATHWAY | 3 |
| REACTOME_CIRCADIAN_CLOCK | 3 |
| REACTOME_APOPTOTIC_EXECUTION_PHASE | 3 |
| WP_IL18_SIGNALING_PATHWAY | 3 |
| WP_FERROPTOSIS | 3 |
| REACTOME_ROS_AND_RNS_PRODUCTION_IN_PHAGOCYTES | 3 |
| WP_PATHWAYS_AFFECTED_IN_ADENOID_CYSTIC_CARCINOMA | 3 |
| REACTOME_FORMATION_OF_FIBRIN_CLOT_CLOTTING_CASCADE | 3 |
| WP_VASOPRESSINREGULATED_WATER_REABSORPTION | 3 |
| REACTOME_G1_S_SPECIFIC_TRANSCRIPTION | 3 |

| | |
|---|---|
| WP_BURN_WOUND_HEALING | 3 |
| WP_NETWORK_MAP_OF_SARSCOV2_SIGNALING_PATHWAY | 4 |
| HALLMARK_INTERFERON_GAMMA_RESPONSE | 4 |
| KEGG_INTESTINAL_IMMUNE_NETWORK_FOR_IGA_PRODUCTION | 4 |
| KEGG_CYTOKINE_CYTOKINE_RECEPTOR_INTERACTION | 4 |
| WP_OVERVIEW_OF_PROINFLAMMATORY_AND_PROFIBROTIC_MEDIATORS | 4 |
| WP_ALLOGRAFT_REJECTION | 4 |
| PID_IL12_2PATHWAY | 4 |
| REACTOME_INTERLEUKIN_10_SIGNALING | 4 |
| REACTOME_CHEMOKINE_RECEPTORS_BIND_CHEMOKINES | 4 |
| KEGG_TYPE_I_DIABETES_MELLITUS | 4 |
| KEGG_AUTOIMMUNE_THYROID_DISEASE | 4 |
| REACTOME_CLASS_A_1_RHODOPSIN_LIKE_RECEPTORS | 4 |
| KEGG_ALLOGRAFT_REJECTION | 4 |
| HALLMARK_INTERFERON_ALPHA_RESPONSE | 4 |
| REACTOME_GPCR_LIGAND_BINDING | 4 |
| WP_TYPE_II_INTERFERON_SIGNALING_IFNG | 4 |
| KEGG_ASTHMA | 4 |
| KEGG_GRAFT_VERSUS_HOST_DISEASE | 4 |
| REACTOME_PEPTIDE_LIGAND_BINDING_RECEPTORS | 4 |
| KEGG_CHEMOKINE_SIGNALING_PATHWAY | 4 |
| WP_SELECTIVE_EXPRESSION_OF_CHEMOKINE_RECEPTORS_DURING_TCELL_POLARIZATION | 4 |
| REACTOME_G_ALPHA_I_SIGNALLING_EVENTS | 4 |
| WP_CHEMOKINE_SIGNALING_PATHWAY | 4 |
| REACTOME_INTERFERON_ALPHA_BETA_SIGNALING | 4 |
| PID_IL27_PATHWAY | 4 |
| PID_IL23_PATHWAY | 4 |
| WP_PROSTAGLANDIN_SIGNALING | 4 |
| PID_IL12_STAT4_PATHWAY | 4 |
| WP_SARSCOV2_INNATE_IMMUNITY_EVASION_AND_CELLSPECIFIC_IMMUNE_RESPONSE | 4 |
| WP_CYTOKINES_AND_INFLAMMATORY_RESPONSE | 4 |
| REACTOME_INTERFERON_SIGNALING | 4 |
| REACTOME_INTERFERON_GAMMA_SIGNALING | 4 |
| KEGG_PRIMARY_IMMUNODEFICIENCY | 5 |

| | |
|---|---|
| KEGG_CELL_ADHESION_MOLECULES_CAMS | 5 |
| WP_TCELL_ACTIVATION_SARSCOV2 | 5 |
| PID_MET_PATHWAY | 5 |
| KEGG_LEISHMANIA_INFECTION | 5 |
| PID_FAK_PATHWAY | 5 |
| REACTOME_GENERATION_OF_SECOND_MESSENGER_MOLECULES | 5 |
| KEGG_NATURAL_KILLER_CELL_MEDIATED_CYTOTOXICITY | 5 |
| REACTOME_COSTIMULATION_BY_THE_CD28_FAMILY | 5 |
| KEGG_T_CELL_RECEPTOR_SIGNALING_PATHWAY | 5 |
| PID_NCADHERIN_PATHWAY | 5 |
| PID_TCR_PATHWAY | 5 |
| WP_MICROGLIA_PATHOGEN_PHAGOCYTOSIS_PATHWAY | 5 |
| WP_MODULATORS_OF_TCR_SIGNALING_AND_T_CELL_ACTIVATION | 5 |
| WP_TCELL_RECEPTOR_TCR_SIGNALING_PATHWAY | 5 |
| REACTOME_PD_1_SIGNALING | 5 |
| WP_ALPHA_6_BETA_4_SIGNALING_PATHWAY | 5 |
| PID_ECADHERIN_STABILIZATION_PATHWAY | 5 |
| PID_CD8_TCR_DOWNSTREAM_PATHWAY | 5 |
| PID_CD8_TCR_PATHWAY | 5 |
| PID_A6B1_A6B4_INTEGRIN_PATHWAY | 5 |
| WP_PATHOGENESIS_OF_SARSCOV2_MEDIATED_BY_NSP9NSP10_COMPLEX | 5 |
| PID_CDC42_PATHWAY | 5 |
| KEGG_B_CELL_RECEPTOR_SIGNALING_PATHWAY | 5 |
| WP_TCELL_ANTIGEN_RECEPTOR_TCR_PATHWAY_DURING_STAPHYLOCOCCUS_AUREUS_INFECTION | 5 |
| WP_REGULATION_OF_ACTIN_CYTOSKELETON | 5 |
| REACTOME_CELL_CELL_COMMUNICATION | 5 |
| KEGG_ANTIGEN_PROCESSING_AND_PRESENTATION | 5 |
| PID_ECADHERIN_NASCENT_AJ_PATHWAY | 5 |
| KEGG_TIGHT_JUNCTION | 5 |
| PID_MTOR_4PATHWAY | 5 |
| PID_NECTIN_PATHWAY | 5 |
| WP_BRAINDERIVED_NEUROTROPHIC_FACTOR_BDNF_SIGNALING_PATHWAY | 5 |
| WP_CANCER_IMMUNOTHERAPY_BY_PD1_BLOCKADE | 5 |
| REACTOME_INTERLEUKIN_2_FAMILY_SIGNALING | 5 |
| WP_B_CELL_RECEPTOR_SIGNALING_PATHWAY | 5 |

| | |
|---|---|
| WP_HEAD_AND_NECK_SQUAMOUS_CELL_CARCINOMA | 5 |
| WP_PI3KAKTMTOR_SIGNALING_PATHWAY_AND_THERAPEUTIC_OPPORTUNITIES | 5 |
| WP_BDNFTRKB_SIGNALING | 5 |
| REACTOME_CELL_JUNCTION_ORGANIZATION | 5 |
| REACTOME_RAF_ACTIVATION | 5 |
| PID_ERBB1_DOWNSTREAM_PATHWAY | 5 |
| REACTOME_RHO_GTPASES_ACTIVATE_NADPH_OXIDASES | 5 |
| PID_IGF1_PATHWAY | 5 |
| WP_EMBRYONIC_STEM_CELL_PLURIPOTENCY_PATHWAYS | 5 |
| WP_IL1_SIGNALING_PATHWAY | 5 |
| KEGG_REGULATION_OF_ACTIN_CYTOSKELETON | 5 |
| PID_EPHB_FWD_PATHWAY | 5 |
| PID_PI3KCI_AKT_PATHWAY | 5 |
| WP_ERBB_SIGNALING_PATHWAY | 5 |
| PID_INSULIN_PATHWAY | 5 |
| WP_EGFEGFR_SIGNALING_PATHWAY | 5 |
| REACTOME_ONCOGENIC_MAPK_SIGNALING | 5 |
| WP_EGFR_TYROSINE_KINASE_INHIBITOR_RESISTANCE | 5 |
| REACTOME_PTK6_REGULATES_RHO_GTPASES_RAS_GTPASE_AND_MAP_KINASES | 5 |
| WP_FRAGILE_X_SYNDROME | 5 |
| REACTOME_GASTRIN_CREB_SIGNALLING_PATHWAY_VIA_PKC_AND_MAPK | 5 |
| WP_HEPATOCYTE_GROWTH_FACTOR_RECEPTOR_SIGNALING | 5 |
| REACTOME_GPVI_MEDIATED_ACTIVATION_CASCADE | 5 |
| WP_LEPTIN_SIGNALING_PATHWAY | 5 |
| REACTOME_INTERLEUKIN_3_INTERLEUKIN_5_AND_GM_CSF_SIGNALING | 5 |
| KEGG_ADHERENS_JUNCTION | 5 |
| WP_GLIOBLASTOMA_SIGNALING_PATHWAYS | 5 |
| WP_SYNAPTIC_SIGNALING_PATHWAYS_ASSOCIATED_WITH_AUTISM_SPECTRUM_DISORDER | 5 |
| WP_GASTRIN_SIGNALING_PATHWAY | 5 |
| REACTOME_ADVANCED_GLYCOSYLATION_ENDPRODUCT_RECEPTOR_SIGNALING | 5 |
| PID_TRKR_PATHWAY | 5 |
| PID_MAPK_TRK_PATHWAY | 5 |

| | |
|---|---|
| REACTOME_THE_ROLE_OF_NEF_IN_HIV_1_REPLICATION_AND_DISEASE_PATHOGENESIS | 5 |
| REACTOME_DNA_REPLICATION | 6 |
| REACTOME_SYNTHESIS_OF_DNA | 6 |
| REACTOME_TNFR2_NON_CANONICAL_NF_KB_PATHWAY | 6 |
| REACTOME_TCR_SIGNALING | 6 |
| REACTOME_APC_C_MEDIATED_DEGRADATION_OF_CELL_CYCLE_PROTEINS | 6 |
| REACTOME_CELL_CYCLE_CHECKPOINTS | 6 |
| REACTOME_APC_C_CDH1_MEDIATED_DEGRADATION_OF_CDC20_AND_OTHER_APC_C_CDH1_TARGETED_PROTEINS_IN_LATE_MITOSIS_EARLY_G1 | 6 |
| REACTOME_SWITCHING_OF_ORIGINS_TO_A_POST_REPLICATIVE_STATE | 6 |
| REACTOME_G2_M_CHECKPOINTS | 6 |
| REACTOME_MITOTIC_G1_PHASE_AND_G1_S_TRANSITION | 6 |
| REACTOME_ORC1_REMOVAL_FROM_CHROMATIN | 6 |
| REACTOME_DNA_REPLICATION_PRE_INITIATION | 6 |
| REACTOME_HOST_INTERACTIONS_OF_HIV_FACTORS | 6 |
| REACTOME_S_PHASE | 6 |
| REACTOME_ANTIGEN_PROCESSING_CROSS_PRESENTATION | 6 |
| REACTOME_SEPARATION_OF_SISTER_CHROMATIDS | 6 |
| REACTOME_G1_S_DNA_DAMAGE_CHECKPOINTS | 6 |
| REACTOME_CDT1_ASSOCIATION_WITH_THE_CDC6_ORC_ORIGIN_COMPLEX | 6 |
| REACTOME_THE_ROLE_OF_GTSE1_IN_G2_M_PROGRESSION_AFTER_G2_CHECKPOINT | 6 |
| REACTOME_STABILIZATION_OF_P53 | 6 |
| REACTOME_NEGATIVE_REGULATION_OF_NOTCH4_SIGNALING | 6 |
| REACTOME_AUF1_HNRNP_D0_BINDS_AND_DESTABILIZES_MRNA | 6 |
| REACTOME_MITOTIC_METAPHASE_AND_ANAPHASE | 6 |
| REACTOME_CROSS_PRESENTATION_OF_SOLUBLE_EXOGENOUS_ANTIGENS_ENDOSOMES | 6 |
| REACTOME_RND3_GTPASE_CYCLE | 7 |
| REACTOME_RHOG_GTPASE_CYCLE | 7 |
| REACTOME_RHOB_GTPASE_CYCLE | 7 |
| REACTOME_RHOJ_GTPASE_CYCLE | 7 |
| REACTOME_RHOV_GTPASE_CYCLE | 7 |
| REACTOME_RND2_GTPASE_CYCLE | 7 |

| | |
|---|---|
| REACTOME_RHOC_GTPASE_CYCLE | 7 |
| REACTOME_RAC3_GTPASE_CYCLE | 7 |
| REACTOME_RHOQ_GTPASE_CYCLE | 7 |
| REACTOME_RHOD_GTPASE_CYCLE | 7 |
| REACTOME_RHOA_GTPASE_CYCLE | 7 |
| REACTOME_RHOU_GTPASE_CYCLE | 7 |
| REACTOME_RND1_GTPASE_CYCLE | 7 |
| REACTOME_RAC1_GTPASE_CYCLE | 7 |
| REACTOME_SIGNALING_BY_EGFR | 8 |
| REACTOME_SIGNALING_BY_ERBB2_IN_CANCER | 8 |
| REACTOME_SHC1_EVENTS_IN_EGFR_SIGNALING | 8 |
| PID_ERBB_NETWORK_PATHWAY | 8 |
| REACTOME_GAB1_SIGNALOSOME | 8 |
| REACTOME_PI3K_EVENTS_IN_ERBB2_SIGNALING | 8 |
| REACTOME_SHC1_EVENTS_IN_ERBB2_SIGNALING | 8 |
| REACTOME_GRB2_EVENTS_IN_ERBB2_SIGNALING | 8 |
| REACTOME_SIGNALING_BY_ERBB2_ECD_MUTANTS | 8 |
| REACTOME_ERBB2_REGULATES_CELL_MOTILITY | 8 |
| REACTOME_ESTROGEN_DEPENDENT_NUCLEAR_EVENTS_DOWNSTREAM_OF_ESR_MEMBRANE_SIGNALING | 8 |
| REACTOME_SIGNALING_BY_FGFR3_FUSIONS_IN_CANCER | 8 |
| PID_ERBB1_INTERNALIZATION_PATHWAY | 8 |
| REACTOME_SIGNALING_BY_ERBB2 | 8 |
| REACTOME_SHC1_EVENTS_IN_ERBB4_SIGNALING | 8 |
| REACTOME_CONSTITUTIVE_SIGNALING_BY_EGFRVIII | 8 |
| WP_ELECTRON_TRANSPORT_CHAIN_OXPHOS_SYSTEM_IN_MITOCHONDRIA | 9 |
| REACTOME_RESPIRATORY_ELECTRON_TRANSPORT_ATP_SYNTHESIS_BY_CHEMIOSMOTIC_COUPLING_AND_HEAT_PRODUCTION_BY_UNCOUPLING_PROTEINS | 9 |
| WP_OXIDATIVE_PHOSPHORYLATION | 9 |
| REACTOME_RESPIRATORY_ELECTRON_TRANSPORT | 9 |
| REACTOME_COMPLEX_I_BIOGENESIS | 9 |
| KEGG_PARKINSONS_DISEASE | 9 |
| KEGG_OXIDATIVE_PHOSPHORYLATION | 9 |
| WP_MITOCHONDRIAL_COMPLEX_I_ASSEMBLY_MODEL_OXPHOS_SYSTEM | 9 |
| REACTOME_INSULIN_RECEPTOR_SIGNALLING_CASCADE | 10 |
| REACTOME_SIGNALING_BY_TYPE_1_INSULIN_LIKE_GROWTH_FACTOR_1_RECEPTOR_IGF1R | 10 |

| | |
|---|---|
| REACTOME_IRS_MEDIATED_SIGNALLING | 10 |
| REACTOME_SIGNALING_BY_INSULIN_RECEPTOR | 10 |
| REACTOME_DOWNSTREAM_SIGNALING_OF_ACTIVATED_FGFR3 | 10 |
| REACTOME_SIGNALING_BY_FGFR1 | 10 |
| REACTOME_DOWNSTREAM_SIGNALING_OF_ACTIVATED_FGFR2 | 10 |
| REACTOME_DOWNSTREAM_SIGNALING_OF_ACTIVATED_FGFR1 | 10 |

**Figures**

**Supplementary Figure 1: IL22 and IL13 producing CD4$^+$ T-cells and iNKT-cells are increased in fistula tissue in cryptoglandular fistulizing disease.** A) The proportion of cytokine-producing CD4$^+$ T-cells in paired peripheral blood and fistula tissue in patients with cryptoglandular fistulizing disease (n=17). B) Representative flow cytometry plot showing production of IL22 and IL13 by CD4$^+$ T-cells in blood and fistula tissue. C) Representative flow cytometry plot showing the division of iNKT-cells into CD8$^+$ CD4$^+$ and CD8$^-$ CD4$^-$ (DN) subsets. D) Frequency of each iNKT-cell subset in blood and fistula tissue in pCD patients. E) Proportion of cytokine producing DN iNKT-cells in blood and fistula samples in cryptoglandular patients. F) Representative flow cytometry plots showing production of IL22 by DN iNKT-cells in blood and fistula. Data expressed as medians. Statistical significance was calculated using Wilcoxon-matched pairs signed rank test. $^{**}P<0.01$ $^{***}P<0.001$, $^{****}P<0.0001$, $^{ns}$not statistically significant.

**Supplementary Figure 2: CD161$^+$ CD4$^+$ T-cells and iNKT-cells are increased in fistula and are superior cytokine producers in pCD.** A) Proportion of CD161 expressing iNKT-cells, CD4$^+$ T-cells, CD8$^+$ T-cells, MAIT cells and γδ T-cells in peripheral blood and fistula tissue samples in pCD (n=14). B) Representative flow cytometry plot showing CD161 expression on $CD4^+$ T-cells and iNKT-cells in blood and fistula tissue. C) Frequency of cytokine producing cells in the $CD161^+$ fraction of $CD4^+$ T-cells compared to the $CD161^-$ fraction. D) Representative flow plots showing production of IL17A, IL22 and TNFα production by $CD161^+$ $CD4^+$ T-cells versus $CD161^-$ $CD4^+$ T-cells. E) Frequency of cytokine producing cells in the $CD161^+$ fraction of DN iNKT-cells compared to the CD161- fraction. F) Representative flow plots showing production of IFNγ, IL13, IL17A and TNFα production by $CD161^+$ DN iNKT-cells versus $CD161^-$ DN iNKT-cells. G) Proportion of $CD4^+$ $CD161^+$ cytokine producing T-cells in blood and fistula tissue. H) Proportion of DN $CD161^+$ cytokine producing iNKT-cells in blood and fistula tissue. Data expressed as medians. Statistical significance was calculated using Wilcoxon-matched pairs signed rank test. *$P<0.05$,**$P<0.01$ ***$P<0.001$, nsnot statistically significant.

**Supplementary Figure 3: $CD161^+$ $CD4^+$ T-cells and iNKT-cells are increased in fistula and are superior cytokine producers in cryptoglandular fistulizing disease.** A) Proportion of CD161 expressing iNKT-cells, $CD4^+$ T-cells, $CD8^+$ T-cells, MAIT cells and γδ T-cells in peripheral blood and fistula tissue samples in cryptoglandular disease (n=17). B) Representative flow cytometry plot showing CD161 expression on CD4+ T-cells and iNKT-cells in blood and fistula tissue. C) Frequency of cytokine producing cells in the $CD161^+$ fraction of $CD4^+$ T-cells compared to the $CD161^-$ fraction. D) Representative flow plots showing production of IL17A, IL22 and TNFα production by $CD161^+$ $CD4^+$ T-cells versus $CD161^-$ $CD4^+$ T-cells. E) Frequency of cytokine producing cells in the $CD161^+$ fraction of DN iNKT-cells compared to the $CD161^-$ fraction. F) Representative flow plots showing production of IFNγ, IL13, IL17A and TNFα production by $CD161^+$ DN iNKT-cells versus $CD161^-$ DN iNKT-cells. G) Proportion of $CD4^+$ $CD161^+$ cytokine producing T-cells in blood and fistula tissue. H) Proportion of DN $CD161^+$ cytokine producing iNKT-cells in blood and fistula tissue. Data expressed as medians. Statistical significance was calculated using Wilcoxon-matched pairs signed rank test. *$P$<0.05, **$P$<0.01 ***$P$<0.001, $^{ns}$not statistically significant.

**Supplementary Figure 4: Transcriptional regulators of T-cell activation and cytokine signalling are activated in fistula tissue and interact with EMT-associated transcription factors.** A) Heatmap of significant differences (FDR<0.05) in transcriptional regulator activity in fistula versus rectal tissue in CD and cryptoglandular fistulizing disease. Positive and negative NES values indicate increased and reduced activity in fistula compared to rectal tissue. B) Network of regulatory interactions among genes in A); edges with triangular and T-shaped ends indicate stimulatory and inhibitory interactions, respectively.

**Supplementary Figure 5: Cytokine signalling, immune activation and wound repair pathways are more activated in CD than cryptoglandular rectal tissue.** A) Distribution of gene expression changes between rectal tissue in donors with CD and cryptoglandular fistulizing disease. B) Hierarchical clustering of pathways regulated in significantly different ways (FDR < 0.05) in rectal tissue from donors with CD and cryptoglandular fistulizing disease. Circular dendrogram shows pathway similarities quantified by Jaccard scores, while inner, middle and outer rings represent assignments to broad functional roles, normalized enrichment scores and false discovery rate, respectively. C) Pathways involved in cytokine signalling, immune activation and wound repair that exhibit altered activity in CD versus cryptoglandular rectal tissue (FDR<0.05).
