## Supplementary figures and images for "Invariant natural killer T-cell and CD4^+^ T-cell derived IL22 is a regulator of epithelial-to-mesenchymal transition and extracellular matrix remodelling in perianal fistulas"

### Supplementary Figure 1

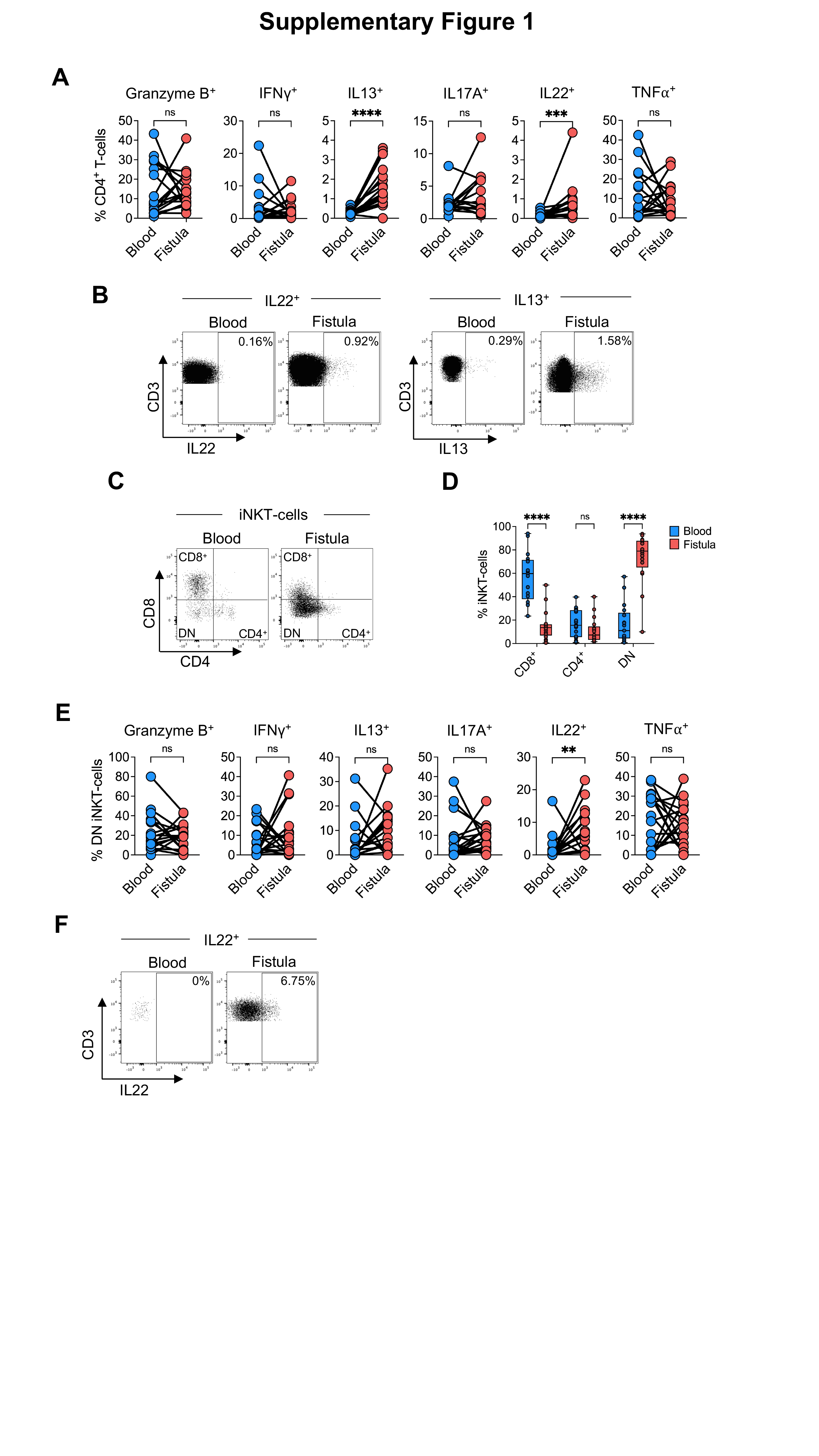

### Supplementary Figure 2

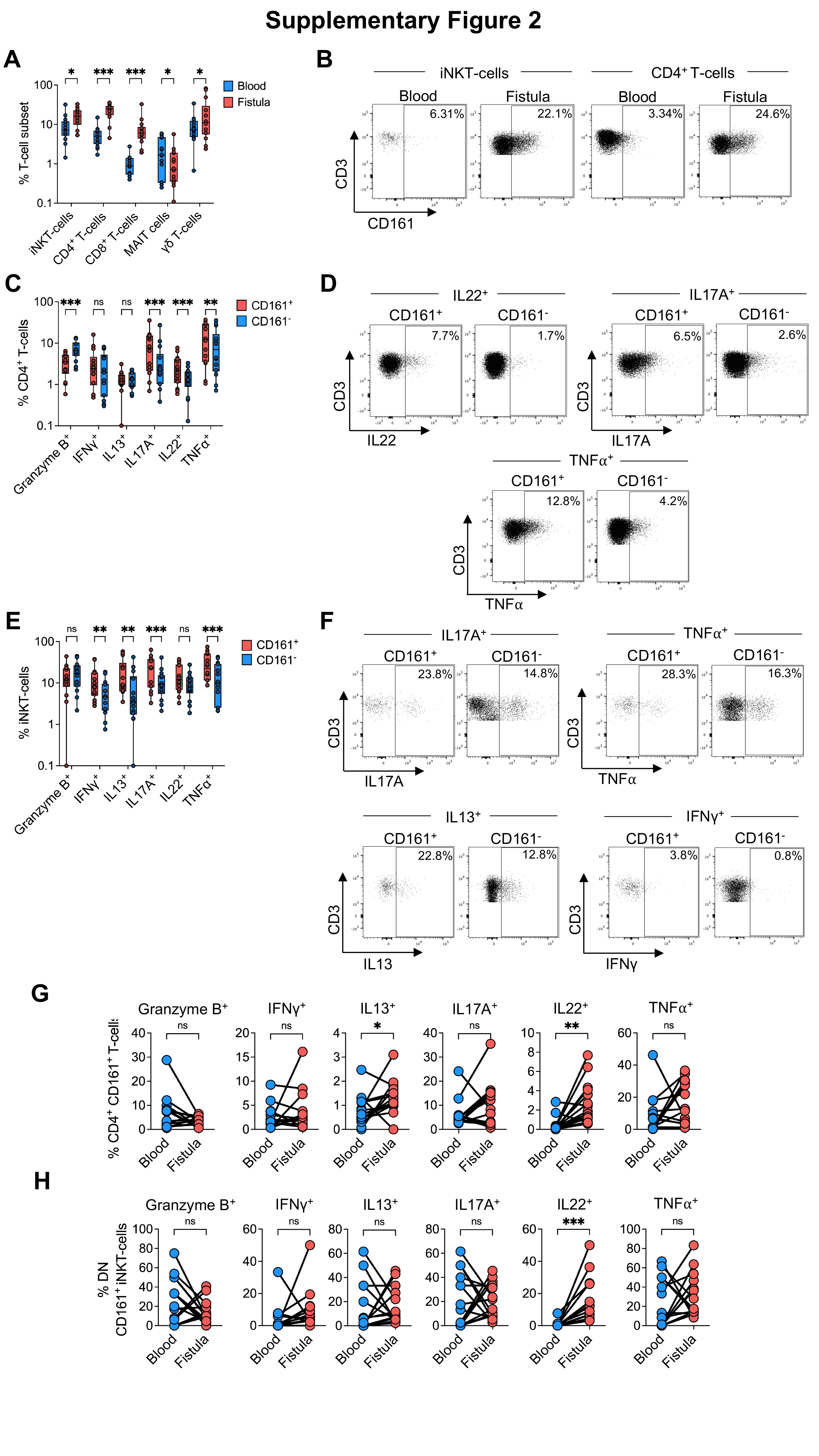

### Supplementary Figure 3

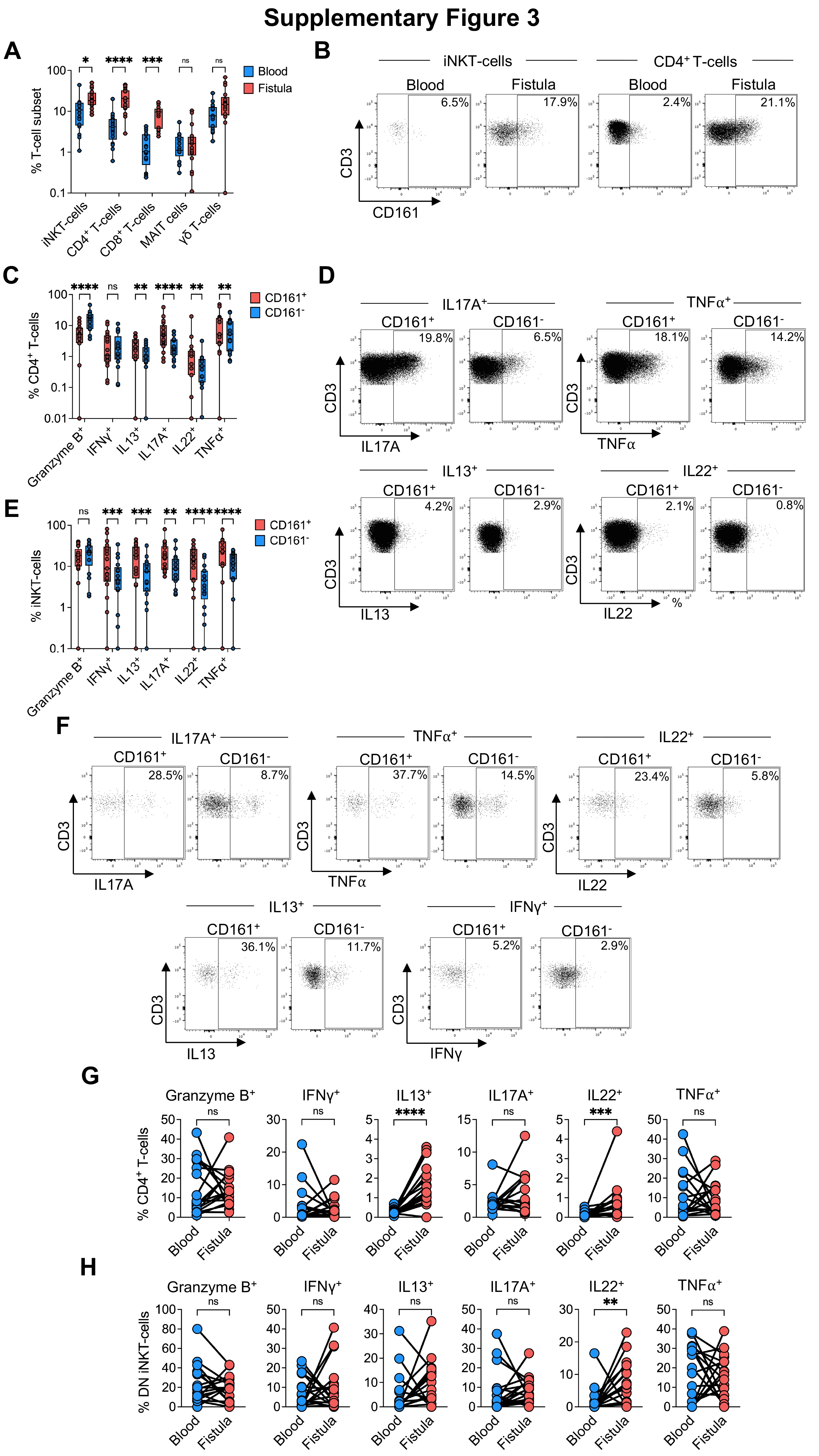

### Supplementary Figure 4

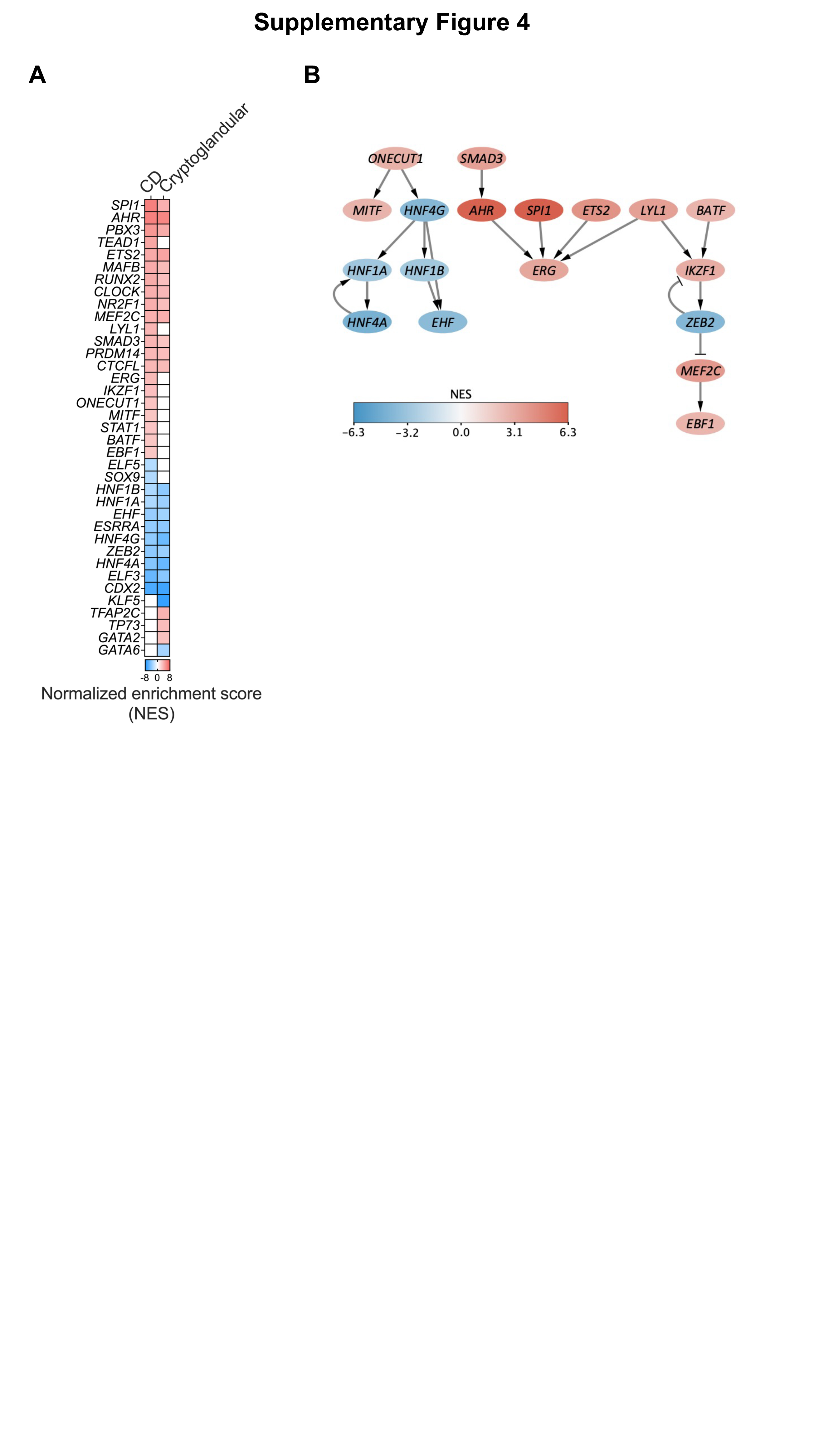

### Supplementary Figure 5

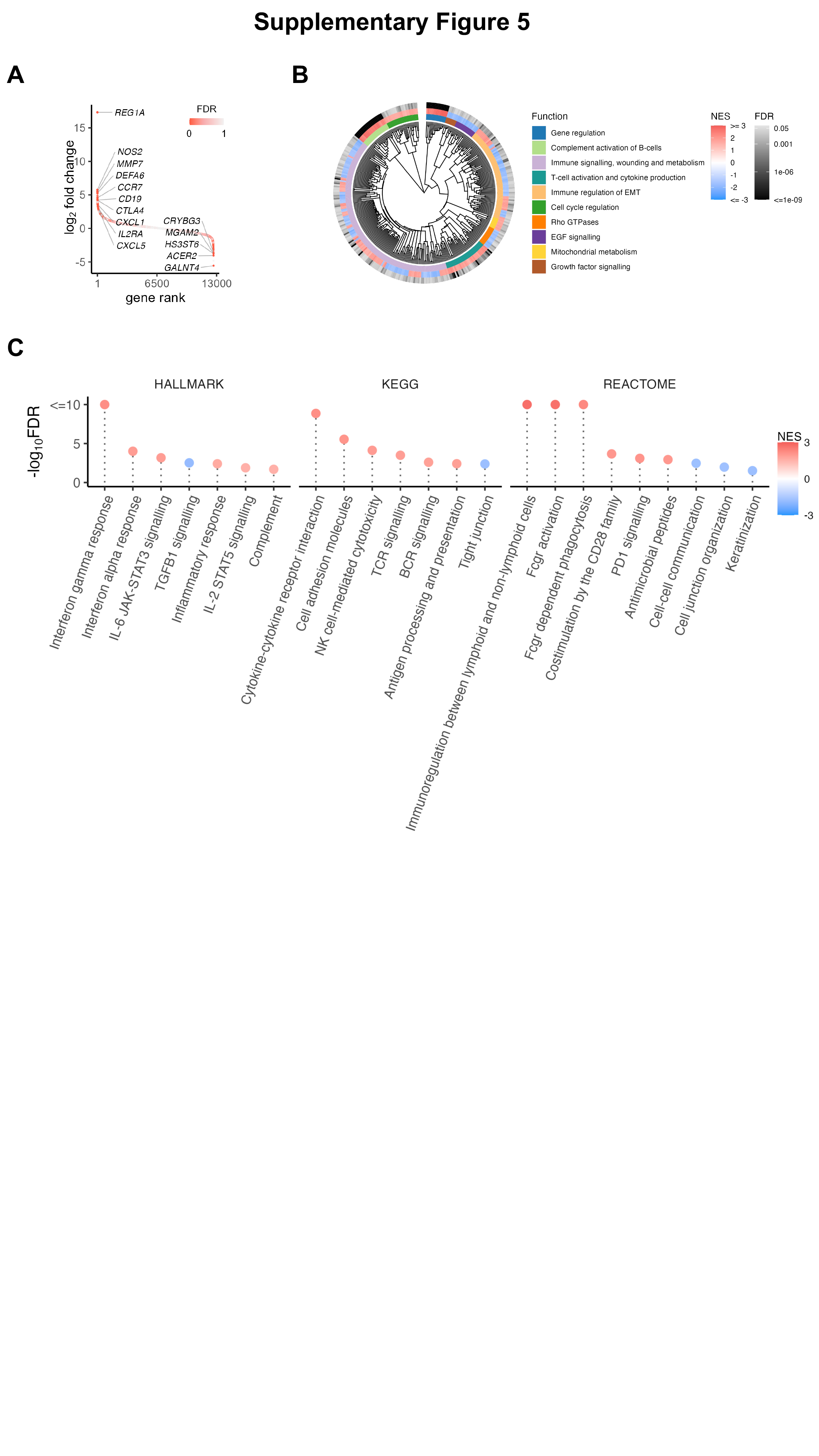
